# Epilepsy and neurobehavioral abnormalities in mice with a *KCNB1* pathogenic variant that alters conducting and non-conducting functions of K_V_2.1

**DOI:** 10.1101/770206

**Authors:** Nicole A. Hawkins, Sunita N. Misra, Manuel Jurado, Nicholas C. Vierra, Kimberly Nguyen, Lisa Wren, Alfred L. George, James S. Trimmer, Jennifer A. Kearney

## Abstract

Developmental and epileptic encephalopathies (DEE) are a group of severe epilepsies that usually present with intractable seizures, developmental delay and are at a higher risk for premature mortality. Numerous genes have been identified as a monogenic cause of DEE, including *KCNB1*. The voltage-gated potassium channel K_V_2.1, encoded by *KCNB1*, is primarily responsible for delayed rectifier potassium currents that are important regulators of excitability in electrically excitable cells, including neurons and cardiomyocytes. The *de novo* pathogenic variant *KCNB1*-p.G379R was identified in an infant with epileptic spasms, atonic, focal and tonic-clonic seizures that were refractory to treatment with standard antiepileptic drugs. Previous work demonstrated deficits in potassium conductance, but did not assess non-conducting functions. To determine if the G379R variant affected clustering at endoplasmic reticulum-plasma membrane junctions K_V_2.1-G379R was expressed in HEK293T cells. K_V_2.1-G379R expression did not induce formation of endoplasmic reticulum-plasma membrane junctions, and co-expression of K_V_2.1-G379R with K_V_2.1-WT lowered induction of these structures relative to K_V_2.1-WT alone, suggesting a dominant negative effect. To model this variant *in vivo*, we introduced *Kcnb1*^G379R^ into mice using CRISPR/Cas9 genome editing. We characterized neurological and neurobehavioral phenotypes of *Kcnb1*^G379R/+^ (*Kcnb1*^R/+^) and *Kcnb1*^G379R/G379R^ (*Kcnb1*^R/R^) mice, and screened for cardiac abnormalities. Immunohistochemistry studies on brains from *Kcnb1*^+/+^ (WT)*, Kcnb1*^R/+^ and *Kcnb1*^R/R^ mice revealed genotype-dependent differences in the levels and subcellular localization of K_V_2.1, with reduced plasma membrane expression of the K_V_2.1-G379R protein, consistent with *in vitro* data. *Kcnb1^R/+^* and *Kcnb1^R/R^* mice displayed profound hyperactivity, repetitive behaviors, impulsivity and reduced anxiety. In addition, both *Kcnb1^R/+^* and *Kcnb1^R/R^* mice exhibited abnormal interictal EEG abnormalities, including isolated spike and slow waves. Spontaneous seizure events were observed in *Kcnb1^R/R^* mice during exposure to novel environments and/or handling, while both *Kcnb1^R/+^* and *Kcnb1^R/R^* mutants were more susceptible to induced seizures. *Kcnb1^R/+^* and *Kcnb1^R/R^* mice exhibited prolonged rate-corrected QT interval on surface ECG recording. Overall, the *Kcnb1^G379R^* mice recapitulate many features observed in individuals with DEE due to pathogenic variants in *KCNB1*. This new mouse model of *KCNB1* associated DEE will be valuable for improving the understanding of the underlying pathophysiology and will provide a valuable tool for the development of therapies to treat this pharmacoresistant DEE.

## Introduction

Developmental and epileptic encephalopathies (DEE) are a group of severe disorders that present early in life and are characterized by paroxysmal activity on EEG, multiple seizure types that are often medically intractable, and serious cognitive, behavioral and neurological deficits (Berg et al., 2010; Scheffer et al., 2017). In 30-50% of infants diagnosed with an epileptic encephalopathy, a causative mutation has been identified in a known epilepsy gene (McTague et al., 2016). Previous work has identified heterozygous *de novo KCNB1* pathogenic variants in individuals with DEE (Allen et al., 2016; Bar et al., 2019; Calhoun et al., 2017; de Kovel et al., 2017; Latypova et al., 2017; Marini et al., 2017; Miao et al., 2017; Saitsu et al., 2015; Thiffault et al., 2015; Torkamani et al., 2014). In addition to seizures, individuals with *KCNB1* variants display comorbidities that include developmental delay, intellectual disability, features of autism spectrum disorder, ADHD and aggression (Bar et al., 2019; Calhoun et al., 2017; de Kovel et al., 2017; Marini et al., 2017). Since the initial reports of *KCNB1* variants in early infantile epileptic encephalopathy type 26 (EIEE26), the phenotype has expanded to include cases with less severe and/or later onset epilepsy or no discrete seizures, and thus the term *KCNB1* encephalopathy encompasses this broader phenotype spectrum.

The *KCNB1*-G379R variant was identified in a male child with seizures commencing at 8 months of age (Torkamani et al., 2014). Glycine 379 is located in the K_V_2.1 channel ion selectivity filter and is an evolutionarily invariant residue critical for potassium selectivity, supporting pathogenicity of the G379R missense variant (Figure 1A) (Kuang et al., 2015). The epilepsy phenotype included infantile spasms, atonic, focal, and tonic-clonic seizures that were refractory to various antiepileptic therapies, including topiramate, valproic acid, pyridoxine, ACTH, and the ketogenic diet. (Torkamani et al., 2014). At 15 months of age, hypsarrhythmia was present on EEG and later in childhood, monitoring revealed multifocal and diffuse polyspikes, polyspike and slow waves, right temporal spike and slow waves, left occipital spikes, and diffuse polyspike bursts (Torkamani et al., 2014). The individual also had severe motor and language delays, autism spectrum disorder, stereotypies, hand-wringing, and strabismus, as well as borderline line QT syndrome (de Kovel et al., 2017; Srivastava et al., 2018; Torkamani et al., 2014).

**Figure 1.**
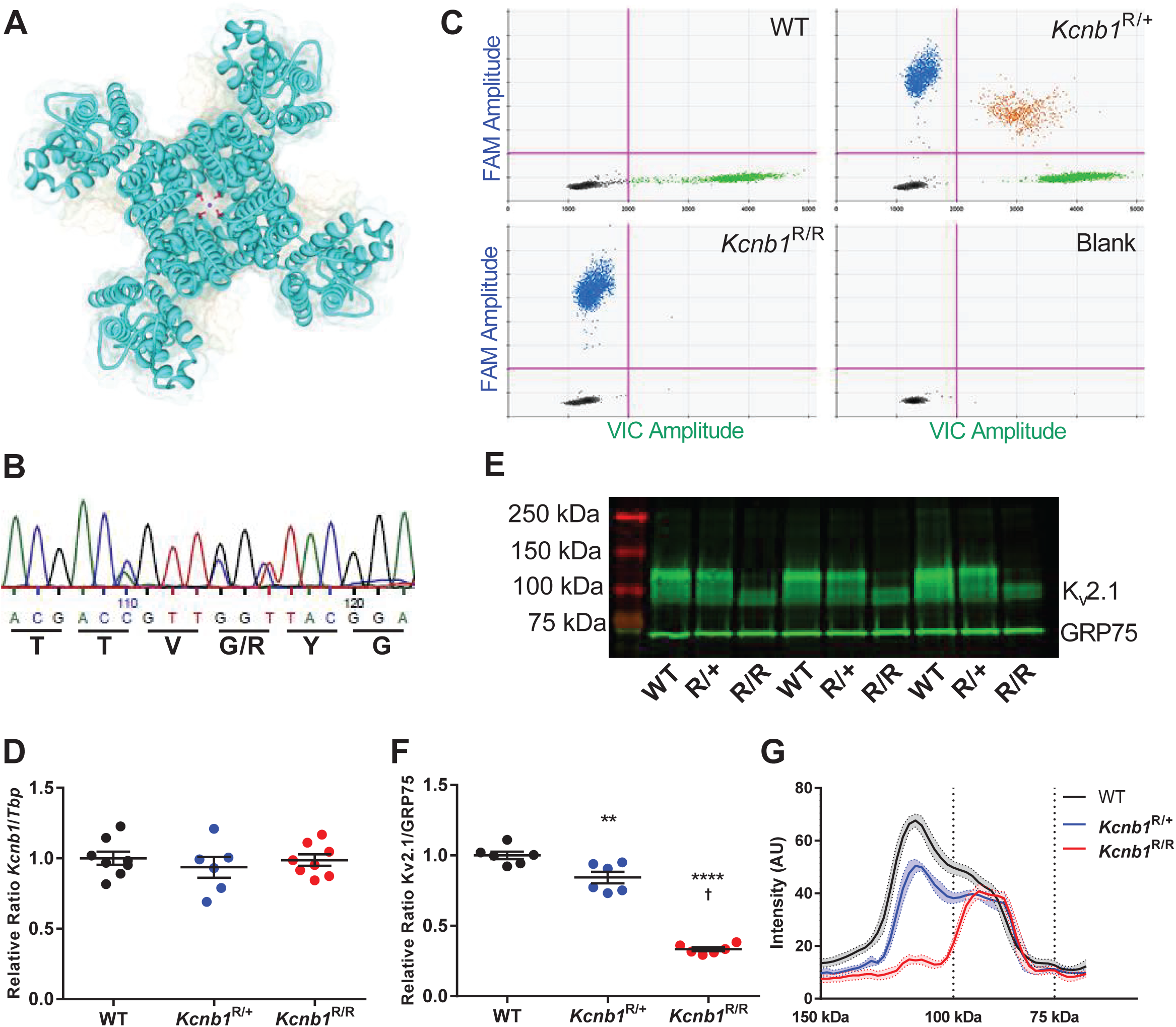
Generation and molecular characterization of *Kcnb1^G379R^* Mice. **A)** Location of glycine-379 in the K_V_2.1 tetrameric potassium channel subunit (PDB 29R9 K_V_2.1/K_V_1.2 chimera). G379 (red) is one of the critical ‘GYG’ residues that determines potassium selectivity and resides in the central pore where backbone carbonyls coordinate the K^+^ ions as they traverse the selectivity filter (black circle). **B)** Sequencing chromatogram of a *Kcnb1* genomic PCR product amplified from a heterozygous *Kcnb1^R/+^* mouse showing heterozygosity for the 3 nucleotides changes introduced by CRISPR/Cas9 editing and homology directed repair. Two nucleotide changes in codon 379 resulted in substitution of arginine for glycine, and a silent substitution in codon 377 disrupted an adjacent PAM site. **C)** Multiplex droplet digital PCR (ddPCR)-based genotyping using hydrolysis probes for the wild-type and G379R alleles distinguishes the three possible genotypes (WT, *Kcnb1^R/+^*, *Kcnb1^R/R^*). Representative two-dimensional scatterplots of each genotype are shown along with a no template control (blank). Each scatterplot represents >10,000 partitioned reactions. Within a scatterplot, each point represents a droplet with a given fluorescence level and droplet colors indicate target amplification about threshold (pink lines). Droplet color code: green=positive for WT; blue=positive for G379R; orange=positive for both; black=negative. **D)** Relative expression of whole brain *Kcnb1* transcript in WT, *Kcnb1^R/+^* and *Kcnb1^R/R^* assays by quantitative RT-ddPCR. There was no difference in transcript expression between genotypes (F(2,19)=0.3806, p>0.7; one-way ANOVA). Symbols represent samples from individual mice and error bars represent SEM with 6-8 mice per genotype. **E)** Representative immunoblots **F)** Quantification of immunoblots showed ≈15% lower K_V_2.1 expression in *Kcnb1^R/+^* and ≈67% lower in *Kcnb1^R/R^* relative to WT (F(2,15)=139.9, p<0.0001; one-way ANOVA). Circles represent samples from individual mice and error bars represent SEM with 6 mice per genotype. **p=0.0049, ****,†p<0.0001, Tukey’s. **G)** Line scan analysis of immunoblots showing a genotype-dependent shift in post-translational state of K_V_2.1 from a heterogeneous pool (≈100-130 kDa) in WT and *Kcnb1^R/+^* samples to approximating the predicted molecular weight of 95 kDa in *Kcnb1^R/R^*. Lines are the average of 6 samples per genotype and SEM is represented by the shading.

*KCNB1* encodes the K_V_2.1 voltage-gated potassium channel subunit that is primarily responsible for delayed rectifier potassium current, an important regulator of neuronal excitability (Guan et al., 2007; Hönigsperger et al., 2017; Liu and Bean, 2014; Murakoshi and Trimmer, 1999; Palacio et al., 2017). K_V_2.1 is expressed at high levels in various neurons and localized to high-density clusters on the soma, proximal dendrites and axon initial segment (Bishop et al., 2015; Du et al., 1998; Sarmiere et al., 2008; Trimmer, 1991). K_V_2.1 clusters align with endoplasmic reticulum (ER) and plasma membrane (PM) junctions, membrane microdomains important in calcium and lipid signaling and homeostasis (Bishop et al., 2018; Bishop et al., 2015; Dickson, 2017; Du et al., 1998; Gallo et al., 2016; Henne et al., 2015; Mandikian et al., 2014; Wu et al., 2017). K_V_2.1 serves a structural role at EM-PM junctions independent of its potassium conducting function by binding to the ER proteins VAPA and VAPB, resulting in the recruitment of the ER to the PM (Fox et al., 2015; Johnson et al., 2018; Kirmiz et al., 2018a; Kirmiz et al., 2018b). In addition to neuronal expression, K_V_2.1 is also expressed in rodent heart within high-density clusters in atrial myocytes, as well as non-clustered K_V_2.1 in ventricular transverse-axial tubules and sarcolemma (O’Connell et al., 2008).

Several *KCNB1* variants have been functionally characterized *in vitro* and shown to exhibit reduced K_V_2.1 conductance, as well as altered ion selectivity, expression and localization (Saitsu et al., 2015; Thiffault et al., 2015; Torkamani et al., 2014). K_V_2.1-G379R channels exhibited loss of voltage-sensitivity and ion-selectivity, resulting in a non-specific cation leak conductance (Torkamani et al., 2014). When co-expressed with wild-type (WT) K_V_2.1 subunits, K_V_2.1-G379R exerted dominant-negative effects on potassium conductance (Torkamani et al., 2014).

In the present study, we evaluated the effect of the DEE-associated *KCNB1*-G379R variant on K_V_2.1 expression and clustering in HEK293T cells and observed a dominant negative effect on ER-PM clustering. To model this variant *in vivo*, we used CRISPR/Cas9 genome editing to generate a mouse model carrying the *KCNB1*-G379R pathogenic variant and characterized Kv2.1 expression and localization, as well as neurological, neurobehavioral, and cardiac phenotypes. *Kcnb1*^G379R^ mice exhibited spontaneous seizures, abnormal EEG patterns, profound hyperactivity, inattention/impulsivity, and reduced anxiety-like behavior. Furthermore, *Kcnb1*^G379R^ mice had prolonged heart-rate corrected QT intervals. These mice recapitulate core features of *KCNB1*-associated encephalopathy and will be a useful tool for understanding disease pathophysiology and evaluating potential therapies.

## Materials and Methods

### HEK293T cell culture, immunocytochemistry, and epifluorescence and TIRF imaging

HEK293T cells were transfected with WT rat K_V_2.1 in pRBG4, rat K_V_2.1 S586A in pCGN, HA-tagged wildtype human K_V_2.1, or HA-tagged WT human K_V_2.1 with the G379R mutation (Kang et al., 2019; Lim et al., 2000; Shi et al., 1994). Cells were transiently transfected using Lipofectamine 2000 following the manufacturer’s protocol within 18 hours of plating. For experiments involving immunocytochemistry, fixation was performed as previously described (Dickson et al., 2016; Kirmiz et al., 2018b). Briefly, HEK293T cells were fixed in 3.2% formaldehyde, blocked and permeabilized. Cells were incubated in monoclonal anti-HA epitope tag antibody 12CA5 (NeuroMab; RRID:AB_ 2532070, pure, 5 μg/mL) and monoclonal anti-VAPA antibody N479/24 (NeuroMab; RRID:AB_ 2722709, 1:5). Coverslips were immunolabeled with Alexa-conjugated fluorescent IgG subclass-specific secondary antibodies (ThermoFisher; 1:1500) and Hoechst 33258 dye (ThermoFisher H1399; 200 ng/mL). Epifluorescence and TIRF imaging of fixed cells and image analysis was performed essentially as described (Bishop et al., 2018; Kirmiz et al., 2018b). Images were obtained with an Andor iXon EMCCD camera installed on a TIRF/widefield equipped Nikon Eclipse Ti microscope using a Nikon LUA4 laser launch with 405, 488, 561 and 647 nm lasers and a 100x PlanApo TIRF/1.49 NA objective run with NIS Elements software (Nikon). Images were collected within NIS Elements as ND2 images. For further details, see supplemental materials and methods.

### Mice

*Kcnb1^G379R/+^* mice on the C57BL/6J inbred strain were generated using CRISPR/Cas9 to introduce the modification of glycine 379 by homology directed repair. Three nucleotide changes were introduced, including two nucleotide changes in codon 379 (GGT > CGC) to maintain similar codon usage, and a silent change to disrupt an adjacent PAM site (C > A) (Figure 1B). A single guide RNA (TAGATGTCTCCGTAACCAA NGG) with good targeting efficiency in Neuro2A cells and low predicted off-target effects, and a 200 bp repair oligo (5’-TCTCCAGCCTGGTCTTCTTTGCCGAGAAGGATGAGGATGACACCAAGTTCAAAAGCA TCCCCGCCTCTTTCTGGTGGGCTACCATCACCATGACGAC**<u>A</u>**GTT**<u>C</u>**G**<u>C</u>**TACGGAGACA TCTACCCTAAGACTCTCCTGGGGAAAATCGTGGGGGGCCTCTGTTGCATTGCCGGTG TCCTGGTGATTGCCCTCCCCATTCCAATT) were microinjected into C57BL/6J embryos by the Northwestern University Transgenic and Targeted Mutagenesis Laboratory.

Potential founders were screened by PCR of genomic DNA using primers outside the repair oligo homology region (Table 1), and the region of interest was TOPO-cloned and Sanger sequenced (n=12-22 clones per founder). The mosaic G379R founder was backcrossed to C57BL/6J mice (Jackson Labs, #000664, Bar Harbor, ME) to generate N1 offspring. N1 offspring were genotyped by Sanger sequencing to confirm transmission of the G379R editing event, and were screened for off-target events by Sanger sequencing of all potential sites with <3 mismatches. N1 mice with the confirmed on-target event and without off-target events were bred with C57BL/6J females to establish the line, *Kcnb1^em1Kea^*, which is maintained by continual backcrossing of heterozygous *Kcnb1^G379R/+^* (*Kcnb1^R/+^*) mice with inbred C57BL/6J mice. Male and female *Kcnb1^R/+^* mice were intercrossed to generate wild-type *Kcnb1^+/+^* (WT), heterozygous *Kcnb1^R/+^*, and homozygous *Kcnb1^R/R^* mice for experiments. For immunohistochemistry experiments, *Kcnb1*^+/-^ and *Kcnb1^-/-^* knockout mice (*Kcnb1^tmDgen^*) that we previously characterized were included for comparison (Speca et al., 2014).

**Table 1.**
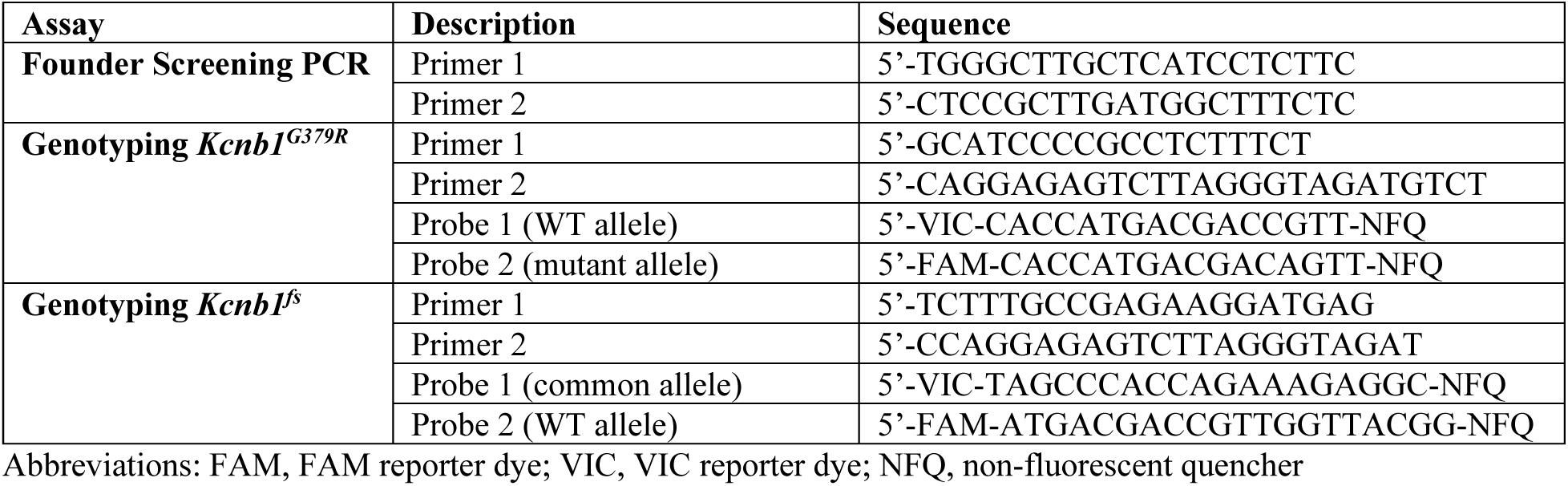
List of Primers and Probes.

Mice were maintained in a Specific Pathogen Free (SPF) barrier facility with a 14-hour light/10-hour dark cycle and access to food and water *ad libitum*. Both female and male mice were used for all experiments. All animal care and experimental procedures were approved by the Northwestern University and UC Davis Animal Care and Use Committees in accordance with the National Institutes of Health Guide for the Care and Use of Laboratory Animals. The principles outlined in the ARRIVE (Animal Research: Reporting of *in vivo* Experiments) guideline and Basel declaration (including the 3 R concept) were considered when planning experiments.

### Genotyping

DNA was isolated from P14 tail biopsies using the Gentra Puregene Mouse Tail Kit according to the manufacturer’s instructions (Qiagen, Valencia, CA). Approximately 250 ng of DNA was digested with BAMH1-HF (R3136, New England Biolabs, Ipswich, MA) at 37°C for 1 hour. Digested DNA was then diluted 1:1 with nuclease-free water and used for template for digital droplet PCR (ddPCR) using ddPCR Supermix for Probes (No dUTP) (Bio-Rad, Hercules, CA, USA) and custom TaqMan SNP Genotyping Assays (Life Technologies, Carlsbad, CA) to detect the mutation (Table 1). Reactions were partitioned into droplets in a QX200 droplet generator (Bio-Rad). PCR conditions were 95°C for 10 minutes, then 44 cycles of 95°C for 30 seconds and 60°C for 1 minute (ramp rate of 2°C/sec) and a final inactivation step of 98°C for 5 minutes. Following amplification, droplets were analyzed with a QX200 droplet reader with QuantaSoft v1.6.6 software (Bio-Rad) (Figure 1C).

### Transcript Analysis

Whole brain RNA was extracted from P27-32 WT, *Kcnb1^R/+^* and *Kcnb1^R/R^* mice. Total RNA was isolated using TRIzol reagent according to the manufacturer’s instructions. First-strand cDNA was synthesized from 2 micrograms of RNA using oligo(dT) primer and Superscript IV reverse transcriptase (RT) according to the manufacturer’s instructions (Life Technologies). First-strand cDNA samples were diluted 1:5 and 5 μl was used as template. Quantitative ddPCR was performed using ddPCR Supermix for Probes (No dUTP) (Bio-Rad) and TaqMan Gene Expression Assays (Life Technologies) for mouse *Kcnb1* (FAM-MGB-Mm00492791_m1) and *Tbp* (VIC-MGB-Mm00446971_m1) as a normalization standard. Reactions were partitioned into a QX200 droplet generator (Bio-Rad). Thermocycling conditions and analysis were performed as described for genotyping. Both assays lacked detectable signal in no-RT and no template controls. Relative transcript levels were expressed as a ratio of *Kcnb1* concentration to *Tbp* concentration, with 6-8 biological replicates per genotype. Statistical comparison between groups was made using ANOVA with Tukey’s post-hoc comparisons (GraphPad Prism, San Diego, CA).

### Immunblotting

Whole brain membrane proteins were isolated from P27-32 WT, *Kcnb1^R/+^* and *Kcnb1^R/R^* mice. Membrane fractions (K_V_2.1: 50 μg/lane; K_V_2.2, AMIGO-1: 100 μg/lane) were separated on a 7.5% SDS-PAGE gel and transferred to nitrocellulose. Blots were probed with monoclonal K_V_2.1 antibody (NeuroMab K89/34; RRID:10673392; 2 μg/mL), monoclonal K_V_2.2 antibody (NeuroMab N372B/1.1; RRID:2315869; 1:2) or monoclonal AMIGO-1 antibody (NeuroMab L86/36.6; RRID: AB_2315799; 1:2) and mouse monoclonal anti-mortalin/GRP75 antibody (NeuroMab N52A/42; RRID:2120479; 1 μg/mL) which served as a normalization control. Alexa Fluor 790 and 680 goat anti-mouse antibodies (Jackson ImmunoResearch, 1:20,000) were used to detect signal on an Odyssey imaging system (Li-COR). Relative protein levels were determined by densitometry with ImageJ (NIH) (Rasband, 1997-2018) or Image Studio (Li-COR) and expressed as a ratio of K_V_2.1, K_V_2.2 or AMIGO-1 to GRP75, with 6-7 biological replicates per genotype. Line scan analysis of K_V_2.1 immunoblots was performed in ImageJ and expressed as the average ± SEM. Statistical comparison between groups was made using one-way ANOVA with Tukey’s post-hoc comparisons (GraphPad Prism).

### Immunohistochemistry

Animals were deeply anesthetized with pentobarbital and transcardially perfused with 4% formaldehyde prepared from paraformaldehyde, in 0.1 M sodium phosphate buffer pH 7.4 (0.1 M PB). Sagittal brain sections (30 µm thick) were prepared and immunolabeled using free-floating methods as previously described (Palacio et al., 2017; Rhodes et al., 2004; Speca et al., 2014). Briefly, free floating sections were blocked and permeabilized. Sections were incubated with primary antibodies and then immunolabeled with Alexa-conjugated fluorescent IgG subclass-specific secondary antibodies and Hoechst 33258 DNA stain (see Table 2 for antibody details). Images were taken using the same exposure time to compare the signal intensity directly. Images were identically processed in Photoshop to maintain consistency between samples. Labeling intensity within *stratum pyramidale* of hippocampal CA1 was measured using a rectangular region of interest (ROI) of 39 μm × 164 μm. To maintain consistency between samples, ROIs were obtained from a region within CA1 near the border of CA1 and CA2. Data points reflect the mean pixel intensity values of this ≈6400 μm^2^ ROI. Values from multiple immunolabels and Hoechst dye were simultaneously measured from the same ROI. Background levels for individual labels were measured from no primary antibody controls and mathematically subtracted from ROI values. For further details, see supplemental materials and methods.

**Table 2.** List of Antibodies.

| **Antibody (Target)** | **Immunogen** | **Manufacturer information** | **Concentration/**<br>**dilution used** | **Figures** |
| --- | --- | --- | --- | --- |
| **N52A/42**<br>**GRP75/mortalin)** | Identified as off-target mAb in screen for anti-SALM2 mAbs | Mouse IgG1 mAb, NeuroMab<br>RRID:AB_2120479 | Pure, 1 µg/mL | 1, 2 |
| **K89/34**<br>**(Kv2.1)** | Synthetic peptide aa 837-853 of rat Kv2.1 | Mouse IgG1 mAb, NeuroMab<br>RRID:AB_RRID:10673392 | Pure | 1 (2 µg/mL);<br>2, 3 (5 µg/mL) |
| **L86/33**<br>**(AMIGO-1)** | Fusion protein aa 394-492 of mouse AMIGO-1 | Mouse IgG2a mAb, NeuroMab<br>RRID:AB_2315798 | Tissue culture supernatant, 1:5 | 2, 3 |
| **Kv2.2C**<br>**(Kv2.2)** | Fusion protein aa 717-907 of rat Kv2.2 long isoform | Rabbit pAb, In-house (Trimmer Laboratory),<br>RRID:AB_2801484 | Affinity purified, 1:100 | 2, 3 |
| **D3/71R**<br>**(Kv2.1)** | Fusion protein aa 506-533 of rat Kv2.1 | Mouse IgG2a recombinant mAb, In-house (Trimmer Laboratory)<br>RRID: AB_2750651 | Tissue culture supernatant, 1:5 | 3 |
| **L80/21**<br>**(Kv2.1)** | Synthetic peptide aa 837-853 of rat Kv2.1 | Mouse IgG3 mAb, NeuroMab<br>RRID:AB_2315862 | Tissue culture supernatant, 1:5 | 3 |
| **L105/31**<br>**(Kv2.1)** | Synthetic peptide aa 596-616 of rat Kv2.1 | Mouse IgG1 mAb, In-house (Trimmer Laboratory)<br>RRID:AB_2801485 | Tissue culture supernatant, 1:5 | 3 |
| **L114/3**<br>**(Parvalbumin)** | Fusion protein amino acids 1-110 of rat Parvalbumin | Mouse IgG2a mAb, NeuroMab<br>RRID:AB_2629420 | Tissue culture supernatant, 1:5 | 3 |
| **L122/6**<br>**(Calretinin)** | Fusion protein amino acids 1-178 of human Calretinin | Mouse IgG2b mAb, NeuroMab<br>RRID:AB_2716256 | Tissue culture supernatant, 1:5 | 3 |
| **12CA5**<br>**(HA epitope)** | Influenza hemagglutinin HA epitope tag | Mouse IgG2b mAb, In-house (Trimmer Laboratory)<br>RRID:AB_2532070 | Pure, 5 µg/mL | 4 |
| **N479/24**<br>**(VAPA)** | Fusion protein aa 1-219 of rat VAPA | Mouse IgG2a mAb, NeuroMab<br>RRID: AB_2722709 | Tissue culture supernatant, 1:5 | 4 |

### Seizure Threshold Testing

Male and female WT, *Kcnb1*^R/+^ and *Kcnb1*^R/R^ mice were tested between 3 and 12 weeks of age by experimenters blinded to genotype. Separate cohorts of mice were used for each inducing agent.

#### Flurothyl seizure induction

Susceptibility to seizures induced by the chemoconvulsant flurothyl (Bis(2,2,2-trifluoroethyl) ether, Sigma-Aldrich, St. Louis, MO) was assessed in WT, *Kcnb1*^R/+^ and *Kcnb1*^R/R^ mice at P70-84. Flurothyl was introduced into a clear, plexiglass chamber by a syringe pump at a rate of 20 μL/min and allowed to volatilize. Latency to first generalized tonic-clonic seizure (GTCS) with loss of posture was recorded (n = 27-32 per genotype). Groups were compared using one-way ANOVA with Tukey’s post-hoc comparisons (GraphPad Prism).

#### 6 Hz seizure induction

Susceptibility to 6 Hz psychomotor seizures was assessed in WT and *Kcnb1*^R/+^ littermates at P19-83. Individual mice were subjected to 6 Hz of auricular stimulation (0.2 ms pulse width, 3 sec duration) using an electroconvulsive unit (Ugo Basile, Gemonio (VA) Italy). Mice that did not exhibit a seizure on the first stimulation were subjected to a second stimulation (≥ 2 mA higher) separated by a delay of ≥15 minutes. Mice were scored for presence or absence of psychomotor seizure activity, defined as forelimb clonus, rearing, paddling or loss of posture. CC_50_ curves are represented with n= 2-10 subjects per current stimulus. Stimulus response curves, CC_50_, and respective confidence intervals (CI) were determined for each sex and genotype using log-probit analysis.

### Video-EEG monitoring

Between P23-25, male and female WT, *Kcnb1*^R/+^ and *Kcnb1*^R/R^ mice were implanted with prefabricated 3-channel EEG headmounts (Pinnacle Technology, Lawrence, KS, USA). Briefly, mice were anesthetized with isoflurane or ketamine/xylazine and placed in a stereotaxic frame. Headmounts with four stainless steel screws that served as cortical surface electrodes were affixed to the skull with glass ionomer cement. Anterior screw electrodes were 0.5–1 mm anterior to bregma and 1 mm lateral from the midline. Posterior screws were 4.5–5 mm posterior to bregma and 1 mm lateral from the midline. EEG1 represents recordings from right posterior to left posterior (interelectrode distance ≈2 mm). EEG2 represents recordings from right anterior to left posterior (interelectrode distance ≈5 mm). The left anterior screw served as the ground connection. Following at least 48 hours of recovery, tethered EEG and video data were continuously collected from freely moving mice Sirenia acquisition software (Pinnacle Technology) as previously described (Hawkins et al., 2016). At least two weeks of EEG was acquired from each subject. Raw data was notch filtered with a 1 Hz window around 60 and 120 Hz prior to analysis. Video-EEG records were analyzed with Persyst13 software (Persyst, Solana Beach, CA, USA), MATLAB (MathWorks, Massachusetts) and EEGLAB (Swartz Center for Computational Neuroscience, California) by a reviewer blinded to genotype. Epileptiform discharges were defined as isolated events with a spike and slow wave morphology, an amplitude of ≥3 times baseline, duration of 150-500 ms, and with increased power in frequencies >20 Hz compared to baseline. Samples with high baseline artifact were excluded from analysis.

### Neurobehavioral Assays

Male and female WT, *Kcnb1*^R/+^ and *Kcnb1*^R/R^ mice were tested between 11 and 15 weeks of age. Male and female mice were tested separately with at least a one-hour delay between sessions. For all experiments, mice were acclimated in the behavior suite with white noise for 1 hour prior to behavioral testing. At the end of each procedure, mice were placed into a clean cage with their original littermates. Behavioral testing was performed by experimenters blinded to genotype. Evaluation occurred over 4 consecutive days: Day 1- neurological exam; Day 2- open field (OF); Day 3- zero maze; and Day 4-cliff avoidance. The marble burying assay was performed on a separate cohort of mice. Statistical comparison between groups were made using one-way ANOVA with Tukey’s post-hoc comparisons for parametric data or two-way repeated measures ANOVA with Sidak’s post-hoc comparisons, unless otherwise indicated.

#### Neurological Exam

Neurological assessment was based on a modified Irwin screen to evaluate baseline behavior of WT and *Kcnb1^G379R^* mice (Irwin, 1968). Individual mice were placed in a small static cage for 3 minutes and observed for transfer behavior, body position, spontaneous activity, tremor, gait, pelvic elevation, tail elevation, palpebral closure, piloerection, air puff startle response, trunk curl, limb grasping, Preyer reflex, provoked salivation, and provoked biting. Each parameter was scored as shown in Table S1 and then summed for a total exam for each mouse. Total exam scores are shown with 19-32 mice per genotype.

#### Open Field

WT and *Kcnb1^G379R^* mice were evaluated for baseline activity in an open field. Individual mice were placed in the center of an open field arena (46 cm × 46 cm) and monitored for 10 minutes. Limelight software (Actimetrics, Wilmette, IL, USA) was used to video record each trial, track the position of the mouse, and calculate distance traveled and relative position in the arena (n= 16-29 mice per genotype).

#### Zero Maze

WT and *Kcnb1^G379R^* mice were evaluated for anxiety-related behavior in an elevated zero maze, a variant on the elevated plus maze that eliminates the ambiguous center region (Shepherd et al., 1994). Individual mice were placed near the closed arm of the maze and allowed to freely explore for 5 minutes. Limelight software was used to video record each trial, track the position of the mouse, and calculate time spent in closed or open arms and total distance traveled (n= 17-28 mice per genotype). Trials where mice jumped off the maze were excluded from the analysis.

#### Cliff Avoidance

WT and *Kcnb1^G379R^* mice were evaluated for impulsive behavior using a cliff avoidance test. Individual mice were placed on an elevated platform (16 cm diameter, 25 cm height) for 7 minutes. Number of peering events (defined as the entire head extending over the edge of the platform) and fall/jumping events (defined as the entire body leaving the platform) were recorded. Jumping events were compared using time-to-event analysis. P-values were determined by LogRank Mantel-Cox tests (n=19-29 per genotype).

#### Marble Burying

Marble movement and burying assay evaluated WT and *Kcnb1^G379R^* mice for phenotypes related to anxiety- and obsessive-compulsive behavior. Individual mice were placed into a static rat cage with 5 cm of corncob bedding and acclimated for 15 minutes. Mice were then briefly removed from the cage while bedding was flattened and 20 marbles were evenly placed across the cage in 5 rows with 4 marbles each, with a small open space at the front of the cage. A baseline image of the cage was taken prior to reintroduction of the mouse and reimaged after the 30-minute trial. The two images were compared for the number of marbles buried, defined by at least ½ of the marble being submerged under the bedding. Total number of marbles moved or buried were compared between groups by two-way repeated measures ANOVA with Sidak’s multiple comparisons test (n= 9-25 mice per genotype).

### Electrocardiography (ECG)

At 8-10 weeks of age, WT, *Kcnb1^R/+^* and *Kcnb^R/R^* mice were anesthetized with isoflurane and surface ECGs were recorded using a digital Dual Bio Amplifier acquisition system and Powerlab software (ADInstruments, Colorado Springs, CO). Baseline ECG was recorded for at least 4 minutes followed by intraperitoneal administration of isoproterenol (1.5 mg/kg) and an additional 10 minutes of recording. Records were analyzed offline using the ECG Module for LabChart software (ADInstruments). Measurement of QT interval was performed on three sequential 10 second intervals at baseline and 6 minutes after the administration of isoproterenol. QT intervals were corrected for heart rate (QT_C_) using Bazett’s formula. Groups were compared using a two-way repeated measures ANOVA test (GraphPad Prism).

### Statistical Analysis

Table 3 summarizes statistical tests used for all comparison along with computed values. Values for post-hoc comparions are reported in the results text and figure legends, and group n’s are reported in figure legends. There were no significant differences between sexes on any measurements except 6 Hz seizure threshold. Thus, groups were collapsed across sex for all variables except the 6 Hz seizure threshold.

**Table 3.**
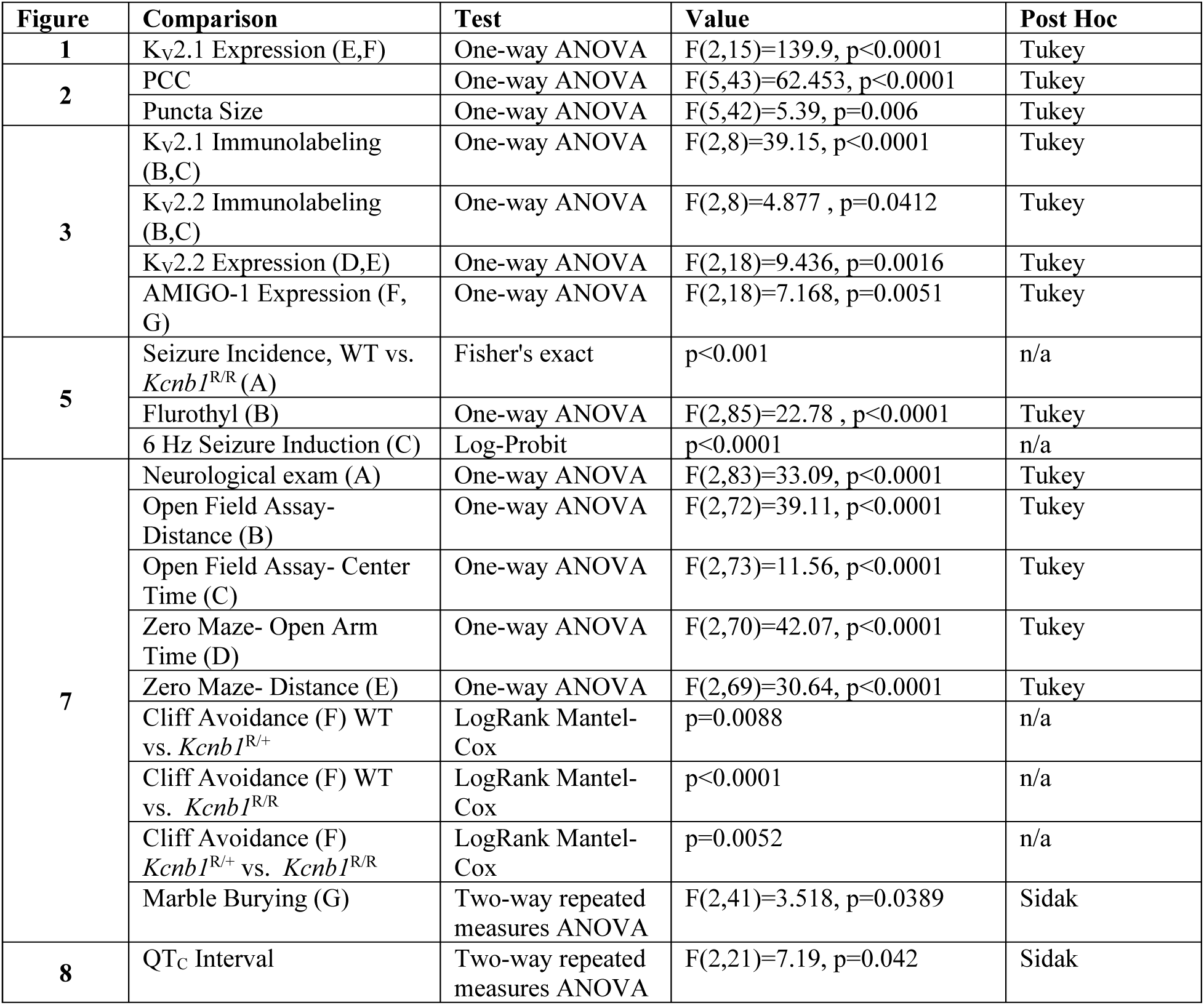
Statistical Comparisons.

## Results

### *KCNB1*-G379R affects subcellular localization of K_V_2.1 in HEK293T cells

Previous reports demonstrated that *KCNB1* variants can alter expression and localization of K_V_2.1 in CHO-K1 and COS-1 cell lines (Thiffault et al., 2015; Torkamani et al., 2014). Additionally, it was shown that exogenously expressed rat K_V_2.1 (rK_V_2.1) in HEK293T cells is specifically localized in large PM clusters similar to those seen in neurons (Bishop et al., 2015; Fox et al., 2015; Johnson et al., 2018; Kirmiz et al., 2018a; Kirmiz et al., 2018b; Mohapatra and Trimmer, 2006; Scannevin et al., 1996; Trimmer, 1991). These clusters are located at ER-PM junctions and result from the interaction of K_V_2.1 with ER proteins VAPA and VAPB (Fox et al., 2015; Johnson et al., 2018; Kirmiz et al., 2018a; Kirmiz et al., 2018b).The clustering of K_V_2.1 and recruitment of VAP proteins at ER-PM junctions is a non-conducting function of K_V_2.1, which instead is dependent upon serine phosphorylation within the PRC domain, a small motif in the K_V_2.1 C-terminus (Johnson et al., 2018; Kirmiz et al., 2018a; Kirmiz et al., 2018b). To determine whether the *KCNB1*-G379R variant affected this non-conducting function of K_V_2.1, we investigated human K_V_2.1 (hK_V_2.1) and hK_V_2.1-G379R expression in HEK293T cells, which lack endogenous K_V_2 channel expression and are of a neuronal lineage (Shaw et al., 2002; Yu and Kerchner, 1998).

We first determined whether hK_V_2.1 formed PM clusters and recruited VAPA proteins to the ER-PM junctions in HEK293T cells. We used two imaging modalities to assess subcellular localization: conventional epifluorescence microscopy to evaluate overall expression pattern, and Total Internal Reflection Fluorescence (TIRF) microscopy to interrogate fluorescent signals within 100-200 nm of the cover slip. The “TIRF field” corresponded to the first ≈1% of the distance into the cell and primarily comprised the PM and PM-associated structures. Expression of hK_V_2.1 in HEK293T cells localized to PM clusters in both epifluorescence and TIRF images, with the majority of hK_V_2.1 present in the clusters (Figure 2A). Expression of hK_V_2.1 led to a significant reorganization in the subcellular localization of VAPA relative to that seen in untransfected cells, such that VAPA was now enriched at K_V_2.1-containing ER-PM junctions (Figure 2A). This colocalization of hK_V_2.1 and VAPA was indicated by a relatively high Pearson’s Correlation Coefficient (PCC) of 0.7 (Figure 2E). Expression of hK_V_2.1-G379R in HEK293T cells suggested a primarily intracellular localization, with lack of concordance between epifluorescence images showing the entire cell and TIRF images showing the region at/near the PM. The hK_V_2.1-G379R PCC value of 0.3 was lower than for hK_V_2.1 (p<0.0001) (Figure 2C,E). Additionally cells expressing hK_V_2.1-G379R had smaller average VAPA cluster size of 0.23±0.02 μm^2^ compared to cells expressing WT hK_V_2.1 (0.63±0.13 μm^2^; p=0.0028, Tukey’s) (Figure 2C,G).

**Figure 2.**
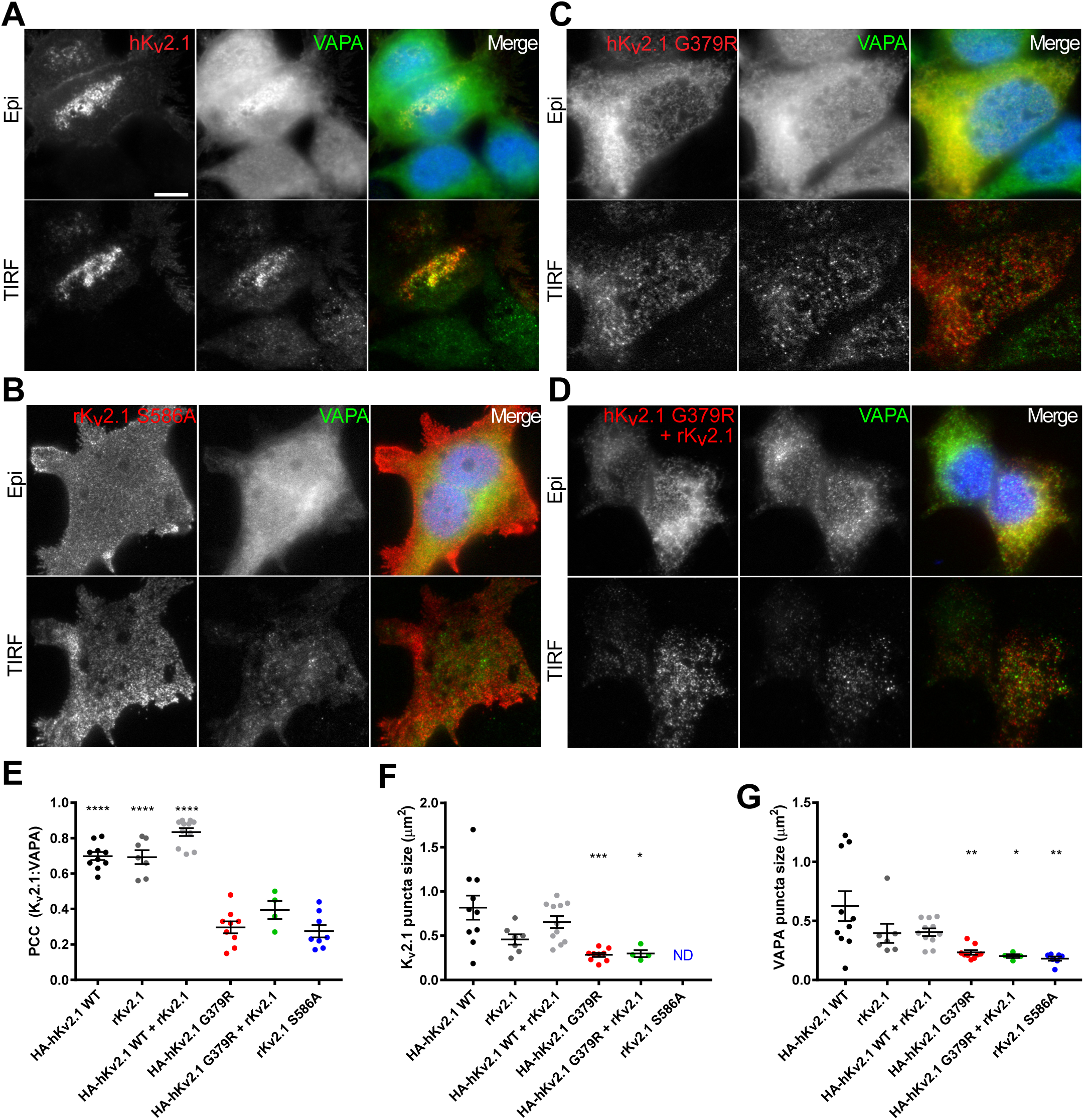
G379R mutation disrupts K_V_2.1-mediated recruitment of VAPA to ER-PM junctions in HEK293T cells. **A-D)** Representative wide-field epifluorescence (“Epi”) and TIRF images of HEK293T cells expressing hK_V_2.1 WT (**A**), rK_V_2.1 S586A (**B**), hK_V_2.1 G379R (**C**) and hK_V_2.1 G379R + rK_V_2.1 (**D**). Red-K_V_2.1, green-VAPA and blue-Hoechst. Scale bar= 10 µm. **E)** Scatter plot of Pearson’s correlation Coefficient (PCC) values for colocalization of K_V_2.1 and VAPA for each transfection condition. PCC values were higher (****p<0.0001, respectively, Tukey’s) for hK_V_2.1 WT: 0.699±0.02, rK_V_2.1: 0.693±0.04 and hK_V_2.1 WT + rK_V_2.1: 0.835±0.02 compared to rK_V_2.1 S586A: 0.275±0.04, hK_V_2.1 G379R: 0.3±0.03 and hK_V_2.1 G379R + rK_V_2.1: 0.4±0.05 (F(5,43)=62.453, p<0.0001; one-way ANOVA). n=4-12 cells. **F)** Scatter plot of K_V_2.1 puncta size for each transfection condition. hK_V_2.1 G379R (0.284±0.2 µm^2^) and hK_V_2.1 G379R + rK_V_2.1 (0.34±0.4 µm^2^) had smaller (p=***0.0006, *p=0.0130, respectively, Tukey’s) K_V_2.1 puncta size compared to hK_V_2.1 WT (0.82±0.14 µm^2^) (F(5,36)=24.59, p<0.0001; one-way ANOVA). n=4-12 cells. ND: Not determined. **G)** Scatter plot of VAPA puncta size for each transfection condition. hK_V_2.1 G379R (0.233±0.02 µm^2^), hK_V_2.1 G379R + rK_V_2.1 (0.203±0.02 µm^2^) and rK_V_2.1 S586A (0.181±0.02 µm^2^) had smaller (**p=0.0028, *p=0.0181, **p=0.0014, respectively, Tukey’s) VAPA puncta size compared to hK_V_2.1 WT (0.626±0.13 µm^2^) (F(5,42)=5.39, p=0.0006; one-way ANOVA). n=4-12 cells. Error bars represent SEM.

The rK_V_2.1-S586A mutation was previously shown to not recruit VAPA to ER-PM junctions (Kirmiz et al., 2018b). As a control, we compared co-localization and VAPA cluster size of rK_V_2.1-S586A with WT hK_V_2.1 or hK_V_2.1-G379R. Consistent with our previous report, expression of rK_V_2.1-S586A resulted in a lower PCC value of 0.28 (p<0.0001) and smaller average VAPA cluster size of 0.18±0.02 μm^2^ compared to WT hK_V_2.1 cluster size of 0.63±0.13 μm^2^ (Figure 2B,E,G). Co-localization PCC values and VAPA cluster sizes were not different between hK_V_2.1 and rK_V_2.1, or hK_V_2.1-G379R and rK_V_2.1-S586A mutants (Figure 2E,G). As an additional control, rK_V_2.1 and hK_V_2.1-G379R were co-expressed and found to have similar localization and size properties of the hK_V_2.1-G379R mutation alone, suggesting G379R may exert a dominant negative effect on co-expressed K_V_2.1 (Figure 2E-G).

### Generation and initial characterization of *Kcnb1^G379R^* mice

To model the DEE-associated *KCNB1*-G379R pathogenic variant *in vivo*, we used CRISPR/Cas9 genome editing to introduce the G379R missense variant in the mouse *Kcnb1* gene. *Kcnb1^G379R/+^* heterozygous and *Kcnb1^G379R/G379R^* homozygous mutants (abbreviated as *Kcnb1^R/+^* and *Kcnb1^R/R^*, respectively) were born at the expected Mendelian ratios. Droplet digital RT-PCR (RT-ddPCR) evaluating whole brain *Kcnb1* transcript indicated no alteration in expression in *Kcnb1*^R/+^ and *Kcnb1*^R/R^ mice relative to *Kcnb1^+/+^* wild-type (WT) littermates, as expected (Figure 1D). We initially observed during the course of routine husbandry that *Kcnb1^G379R^* mice exhibited seizures following handling, overt home cage hyperactivity, repetitive jumping, and occasional unexpected deaths in otherwise healthy appearing animals (median age at death P27, range P21-172).

### Lower K_V_2.1 channel expression in *Kcnb1^G379R^* mice

To evaluate the effect of the *Kcnb1^G379R^* variant on K_V_2.1 protein expression, we first performed immunoblotting using whole brain membrane preparations. These analyses revealed that whole brain expression of K_V_2.1 was ≈15% lower in *Kcnb1^R/+^* heterozygotes and 67% lower in *Kcnb1^R/R^* homozygous mice compared to WT littermates (Figure 1E,F). There was also an evident shift in the relative electrophoretic mobility (M_r_) from the fully post-translationally modified form (≈125 kDa) toward an M_r_ of ≈96 kDa, suggesting a deficit in the processing of the *Kcnb1^G379R^* variant into the the mature, fully post-translationally modified species (Figure 1E,G) (Misonou et al., 2004; Murakoshi et al., 1997).

To assess the impact of the *Kcnb1^G379R^* variant on neuroanatomy and K_V_2.1 expression, we performed immunolabeling of brain sections from WT, *Kcnb1^R/+^* and *Kcnb1^R/R^* mice, as well as *Kcnb1^+/-^* and *Kcnb1^-/-^* mice for comparison (Speca et al., 2014). Based on labeling with the DNA-specific dye Hoechst 33258 (data not shown) and consistent with previous observations in *Kcnb1*^-/-^ mice (Figure 3A), the *Kcnb1^G379R^* variant had no overt effect on gross hippocampal neuroanatomy (Speca et al., 2014). Densitometry of fluorescent signal intensity revealed lower K_V_2.1 immunolabeling (green) in hippocampal, subicular and neocortical neurons from *Kcnb1*^G379R^ mice compared to WT (Figure 3B,C), with ≈55% less K_V_2.1 labeling in *Kcnb1*^R/+^ and ≈80% less in *Kcnb1*^R/R^ relative to WT (p=0.0009 and p<0.0001, respectively). However, the lower K_V_2.1 immunolabeling in *Kcnb1^R/R^* mice was not as dramatic as the complete lack of detectable K_V_2.1 observed in *Kcnb1^-/-^* mice (Figure 3B,C).

**Figure 3.**
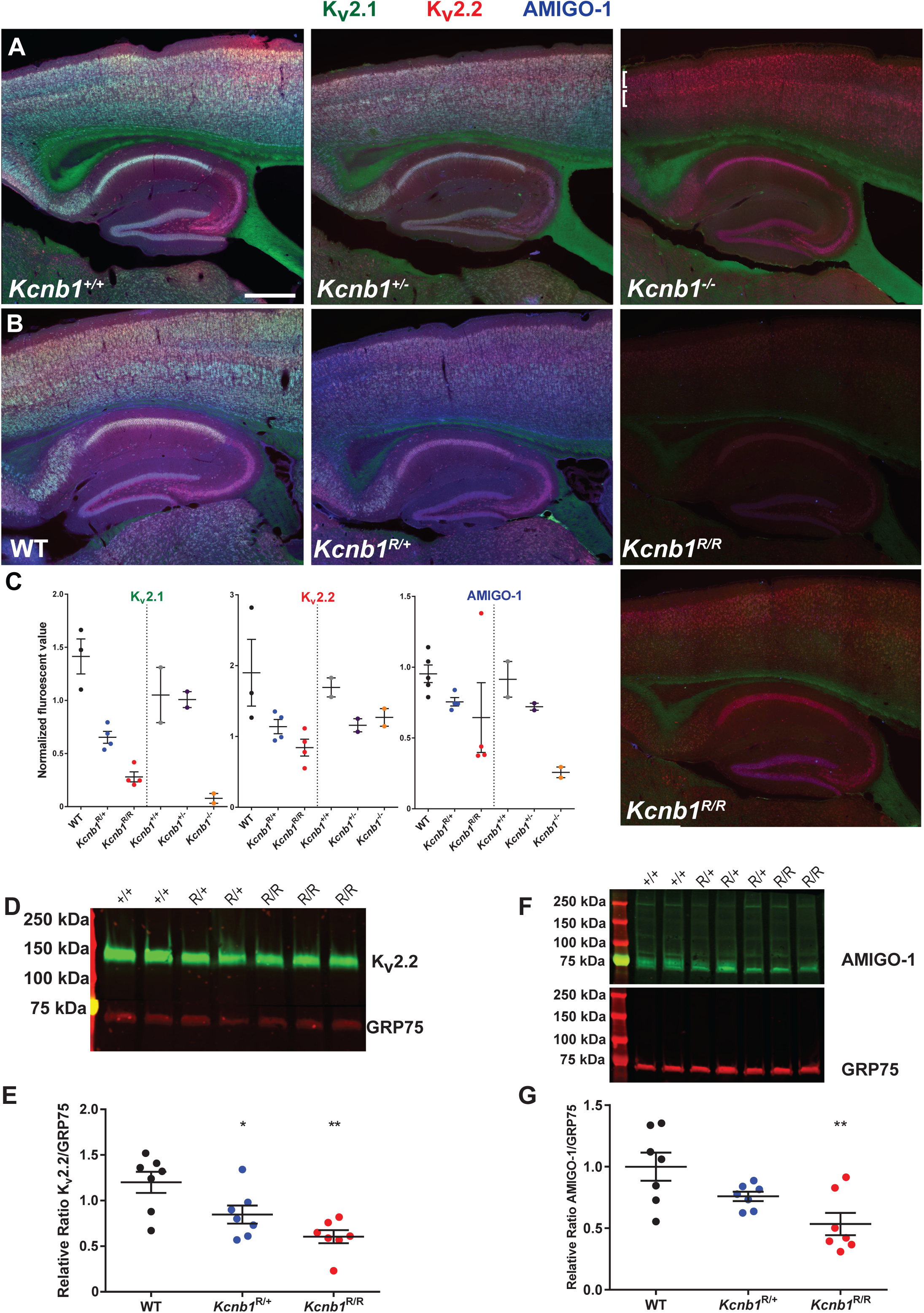
Immunolabeling of *Kcnb1^G379R^* brain sections and membrane preps. **A)** Representative images of somatosensory cortex and hippocampus from littermate *Kcnb1*^+/+^, *Kcnb1*^+/-^ and *Kcnb1*^/-^ mice. While K_V_2.1 signal is lost, there is persistent overlap of K_V_2.2 and AMIGO-1 in K_V_2.2-positive neurons in hippocampus and cortical layers 2 and 5a (brackets, magenta). Green-K_V_2.1, red-K_V_2.2 and blue-AMIGO-1.Scale bar = 500 μm **B)** Representative images of somatosensory cortex and hippocampus from littermate WT, *Kcnb1*^R/+^ and *Kcnb1*^R/R^ mice. Signal intensities for K_V_2.1, K_V_2.2, and AMIGO-1 immunolabeling are lower in *Kcnb1*^R/R^ mice. Bottom right *Kcnb1^R/R^* panel shows that when the intensity of enhances, the remaining signal is predominantly K_V_2.2 and AMIGO-1 (magenta). **C)** Summary graphs of normalized mean fluorescence intensity from a region of interest within *stratum pyramidale* of CA1. K_V_2.1: F(2,8)=39.15; p<0.0001; one-way ANOVA. †p=0.034 (*Kcnb1*^R/+^:*Kcnb1*^R/R^), ***p=0.0009, ****p<0.0001. K_V_2.2: F(2,8)=4.877; p=0.0412; one-way ANOVA. *p=0.0364. **D)** Representative K_V_2.2 immunoblot **E)** Quantification of K_V_2.2 immunoblots showed ≈30% lower expression in *Kcnb1^R/+^* and ≈50% lower in *Kcnb1^R/R^* relative to WT (F(2,18)=9.436, p=0.0016; one-way ANOVA). Circles represent samples from individual mice and error bars represent SEM with 7 mice per genotype. *p<0.049, **p<0.001, Tukey’s. **F)** Representative AMIGO-1 immunoblot **G)** Quantification of AMIGO-1 immunoblots showed ∼25% lower expression in *Kcnb1^R/+^* and ≈50% lower in *Kcnb1^R/R^* relative to WT (F(2,18)=7.168, p=0.0051; one-way ANOVA). Circles represent samples from individual mice and error bars represent SEM with 7 mice per genotype. **p=0.0037, Tukey’s.

Additional validation of lower K_V_2.1 expression due to the *Kcnb1^G379R^* variant was performed by immunolabeling with three additional K_V_2.1 antibodies, D3/71R, L105/31 and L80/21, each of which detected a similar reduction in K_V_2.1 (Figure 4A). Both D3/71 and L105/31 have binding sites on the K_V_2.1 protein distinct from one another and from the other anti-K_V_2.1 antibodies used (Table 2). Immunolabeling with antibodies against three different regions of K_V_2.1 supports that the *Kcnb1^G379R^* variant yields lower K_V_2.1 expression relative to WT. Importantly, to control for possible technical artifacts as the basis of the differing degrees of immunolabeling, parvalbumin and calretinin immunolabeling was performed and found to not differ between the brain sections from *Kcnb1*^R/+^, *Kcnb1*^R/R^, and WT mice (Figure 4B).

**Figure 4.**
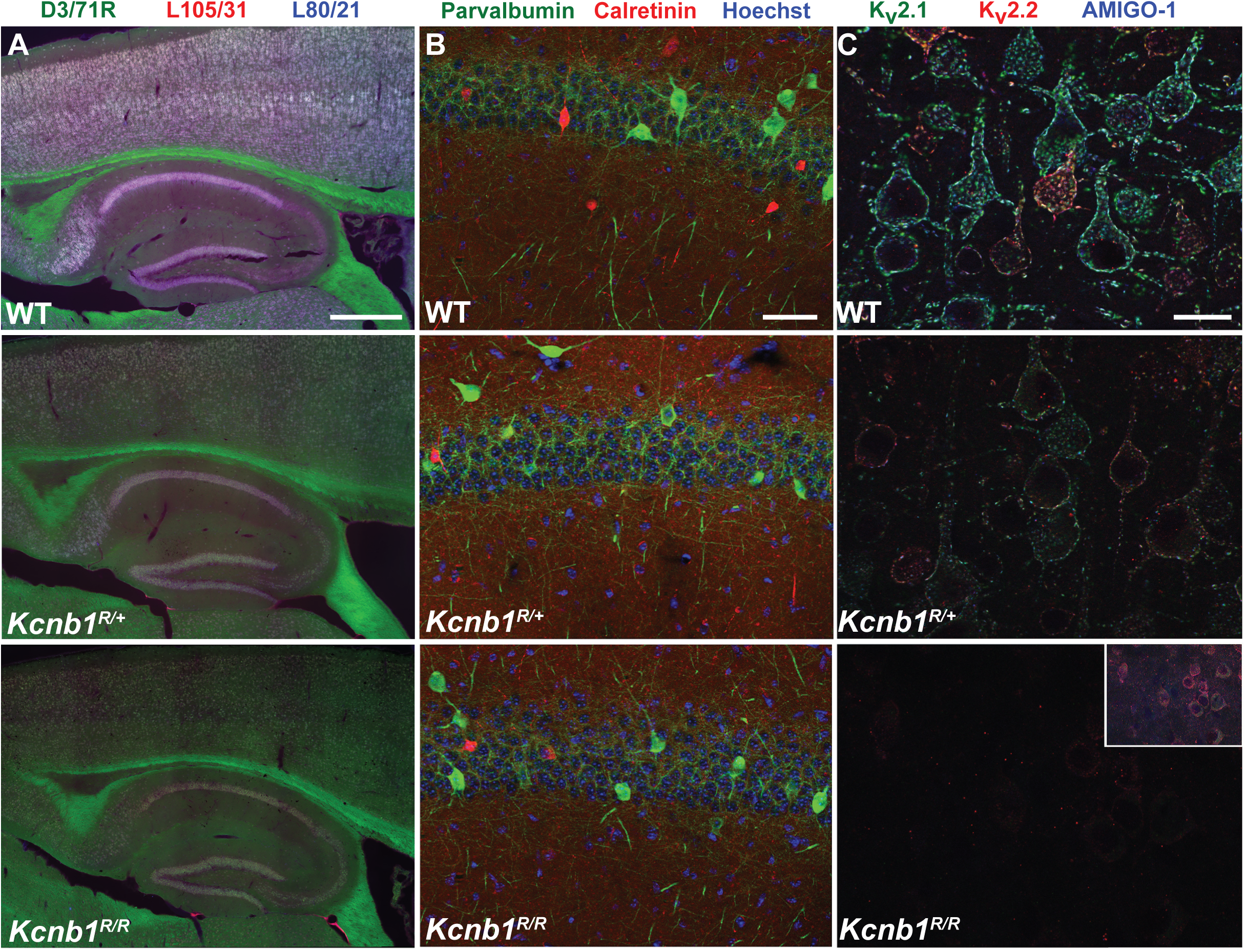
Immunolabeling in *Kcnb1^G379R^* brain sections. **A)** Representative images of somatosensory cortex and hippocampus from littermate WT, *Kcnb1*^R/+^ and *Kcnb1*^R/R^ mice. Immunolabeling with three distinct K_V_2.1 antibodies show distinct K_V_2.1 loss in *Kcnb1^R/+^* and *Kcnb1^R/R^* mice. Green-D3/71R, red-L105/31 and blue-L80/21. Scale bar = 500 μm. **B)** Representative images of CA1 from littermate WT, *Kcnb1*^R/+^ and *Kcnb1*^R/R^ mice. No genotype-specific differences in immunolabeling are apparent. Green-parvalbumin, red-calretinin and blue-Hoechst 33258. Scale bar = 50 μm **C)** Representative high magnification images of layer 5b of somatosensory cortex from littermate WT, *Kcnb1*^R/+^ and *Kcnb1*^R/R^ mice. Inset panel is 3x reduced image to show immunolabeling of *Kcnb1*^R/R^ mice is predominantly intracellular compared to WT and *Kcnb1*^R/+^ mice. Green-K_V_2.1, red-K_V_2.2 and blue-AMIGO-1. Scale bar = 20 μm.

*Kcnb1^G379R^* also affects expression of K_V_2.2 and the K_V_2 channel auxiliary subunit AMIGO-1. Immunolabeling experiments performed on WT, *Kcnb1*^R/+^ and *Kcnb1*^R/R^ brain sections revealed lower K_V_2.2 expression in *Kcnb1*^R/R^ versus WT mice (Figure 3B,C) (p<0.036). To further evaluate the effect of the *Kcnb1^G379R^* variant on K_V_2.2 and AMIGO-1 protein expression, immunoblotting was performed on whole brain membrane preparations (Figure 3D,E). Immunoblotting showed that whole brain expression of K_V_2.2 was ≈30% lower in *Kcnb1*^R/+^ and ≈50% lower in *Kcnb1*^R/R^ mice compared to WT littermates. Immunoblotting for AMIGO-1 whole brain expression revealed ≈25% lower expression in *Kcnb1*^R/+^ and ≈50% lower expression in *Kcnb1*^R/R^ mice compared to WT littermates (Figure 3F,G). The observed reduction of AMIGO-1 in *Kcnb1^G379R^* mice is in contrast to our previous studies of brain sections from *Kcnb1^+/-^* and *Kcnb1^-/-^* mice (Bishop et al., 2018), in which AMIGO-1 immunolabeling was only lost in neurons that typically express high levels of K_V_2.1 relative to K_V_2.2, but was expressed normally in neurons that express K_V_2.2, such as CA1 pyramidal neurons (Figure 3A).

### *Kcnb1*^G379R^ affects K_V_2.1 subcellular localization

High magnification images of pyramidal neurons in layer 5b of the cortex depict that the *Kcnb1^G379R^* variant affects subcellular localization of K_V_2.1 channels (Figure 4C). *Kcnb1*^G379R^ mutant channels have a primarily intracellular distribution with a reduced intensity consistent with immunoblot results (Figure 1E,F, 4C). Moreover, the intracellular accumulation of residual K_V_2.1 identified in *Kcnb1^R/R^* mice is consistent with the shift in M_r_ to a species of ≈96 kDa (Figure 1E,G). This suggests *Kcnb1*^G379R^ mutant channels fail to reach the PM, and also do not acquire post-translational modifications, characteristic of the fully mature K_V_2.1 channel (Misonou et al., 2004; Murakoshi et al., 1997).

### Handling-induced seizures in *Kcnb1^G379R^* mice

Based on initial observations of seizures during the course of routine husbandry, we systematically assessed spontaneous behavioral seizures by individually transferring mice to clean cages and observing for 1 min once per week between 6 and 12 weeks of age. During the course of those observations, 42% (10 of 24) *Kcnb1^R/R^* mice exhibited behavioral seizures with forelimb clonus, rearing and falling, sometimes followed by wild running (Figure 5A, Supplemental Video S1). In contrast, *Kcnb1^R/+^* heterozygous mice (n= 23) and WT littermates (n= 23) did not exhibit any seizures during these observations (p<0.001). Rarely, similar behavioral seizures were observed in *Kcnb1^R/+^* heterozygotes, but at older ages (>4 months).

**Figure 5.**
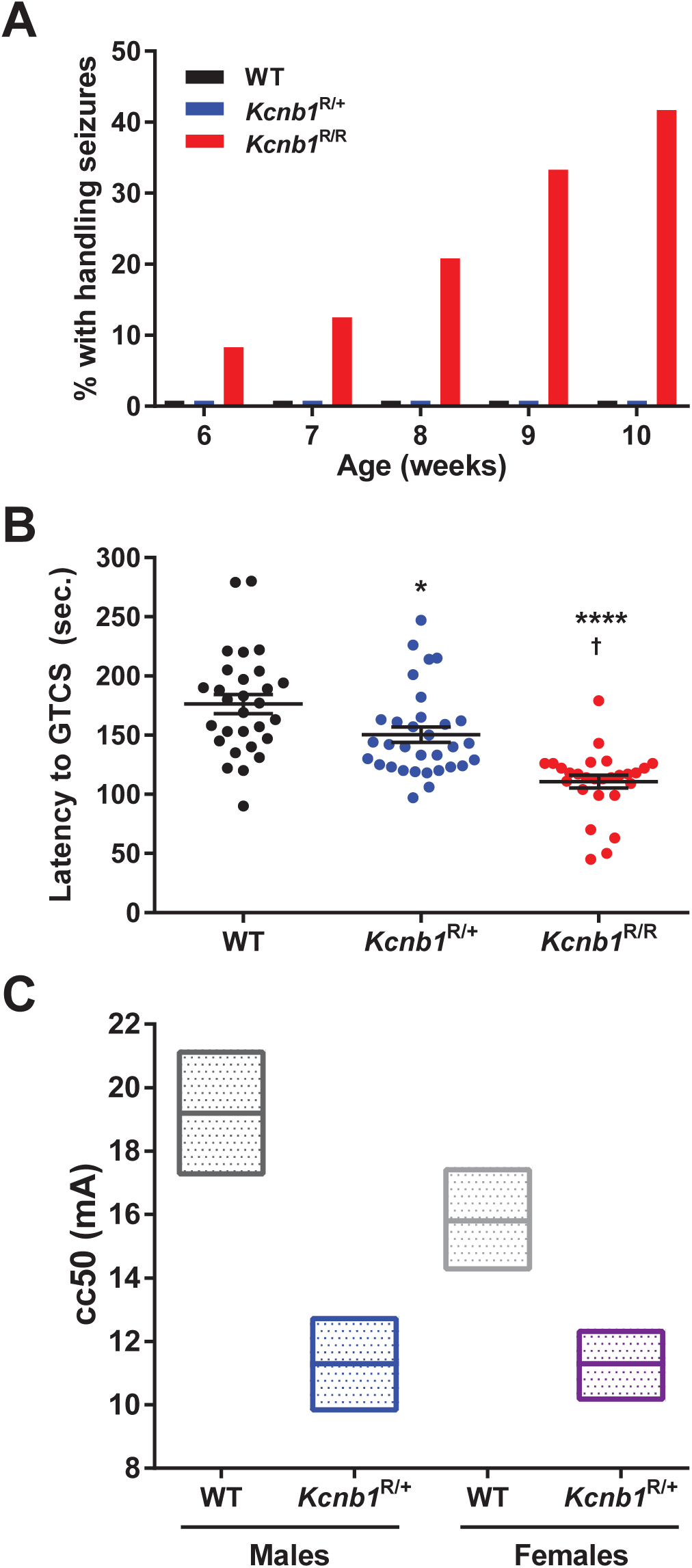
*Kcnb1^R/+^* and *Kcnb1^R/R^* mice are sensitive to induced seizures. **A)** Handling-induced seizures were observed in *Kcnb1^R/R^* mice (n=24) at 6-12 weeks of age, while they were not observed in *Kcnb1^R/+^* (n=23) or WT (n=23) mice in this age window (p<0.001, WT vs *Kcnb1^R/R^*, Fisher’s Exact). Rare handling-induced seizures were observed in *Kcnb1^R/+^* mice at >4 months of age. **B)** Latency to flurothyl-induced GTCS assessed at P70-84 was affected by genotype (F(2,85)=22.78, p<0.0001, one-way ANOVA). *Kcnb1^R/+^* mice had a reduced seizure threshold compared to WT with an average latency of 150 ± 6 and 176 ± 8 sec, respectively (*p=0.0187, Tukey’s). *Kcnb1^R/R^* mice had an average latency of 111 ± 5 sec, which was lower compared to WT (****p<0.0001, Tukey’s) and *Kcnb1^R/+^* (†p= 0.0002) with 27-32 mice per genotype. Error bars represent SEM. **C)** Susceptibility to psychomotor seizures induced by 6 Hz stimulation was assessed at P19-83. *Kcnb1*^R/+^ mice had lower CC_50_ values (convulsive current, 50% with seizures) compared to WT (p<0.0001, log-probit). The CC_50_ (95% confidence interval) of male WT and *Kcnb1^R/+^* mice was 19.2 mA (17.2 to 21.2) and 11.3 mA (9.8 to 12.8), respectively. The CC_50_ of female WT and *Kcnb1^R/+^* mice was 15.8 mA (14.2 to 17.5) and 11.3 mA (10.1 to 12.4), respectively. Floating box graph represents 95% CI and CC_50_ (line). CC_50_ and CI were determined using log-probit analysis with n= 2-10 per current.

### Reduced threshold for induced seizures in *Kcnb1^G379R^* mice

We investigated whether the *Kcnb1^G379R^* mutation influenced latency to seizures induced by the volatile chemoconvulsant flurothyl, a GABA_A_ antagonist (van Vliet, 2017). Latency to the first generalized tonic-clonic seizure with loss of posture was lower in *Kcnb1*^R/R^ mice compared to WT (p<0.0001) and *Kcnb1^R/+^* mice (p=0.0002). *Kcnb1*^R/+^ mice also had a lower threshold for flurothyl-induced seizures than WT mice (p=0.0187). Average latency was 111 ± 5 sec for *Kcnb1^R/R^* mice, 150 ± 6 sec for *Kcnb1^R/+^* mice, and 176 ± 8 sec for WT littermates. (Figure 5B).

We next investigated whether the *Kcnb1* mutation influenced latency to psychomotor seizures induced by long duration, low frequency stimulation (6 Hz, 3 sec), a model of pharmacoresistant focal seizures (van Vliet, 2017). *Kcnb1^R/+^* heterozygotes and WT littermates were subjected to 6 Hz stimulation and scored for the presence or absence of seizure activity (forelimb clonus, rearing, paddling or loss of posture). Convulsive current (CC) curves were generated as previously described (Finney, 1971), and population CC_50_ values (CC at which 50% of mice seized) were determined for each genotype and sex. Both male and female *Kcnb1*^R/+^ mice had lower CC_50_ values compared to sex-matched WT controls (p<0.0001) (Figure 5C). The CC_50_ values (95% confidence interval) for WT and *Kcnb1*^R/+^ male mice were 19.2 mA (17.2 to 21.2) and 11.3 mA (9.8 to 12.8), respectively. The CC_50_ values for WT and *Kcnb1*^R/+^ female mice were 15.8 mA (14.2 to 17.5) and 11.3 mA (10.1 to 12.4), respectively. The relative genotype-dependent shift was similar for males and females.

### EEG abnormalities in *Kcnb1^G379R^* mice

The handling-induced behavioral seizure events described above corresponded with electrographic generalized tonic-clonic seizures detected during video-EEG monitoring of *Kcnb1^R/+^*, *Kcnb1^R/R^* and WT mice at 3 to 6 weeks of age (Figure 6). In addition, several types of interictal EEG abnormalities were observed. Both *Kcnb1^R/+^* and *Kcnb1^R/R^* mice exhibited isolated spike and slow wave complexes (Figure 6B-E) that have an increase in power across low and high frequencies up to 170 Hz (Figure 6E). The occurrence of the isolated spike and slow wave complexes was elevated relative to WT littermates (Figure 6F) when quantified over a 24-hour period. In addition, homozygous *Kcnb1^R/R^* mice displayed recurrent, brief runs of rhythmic slow spike and wave complexes (1-2 Hz).

**Figure 6.**
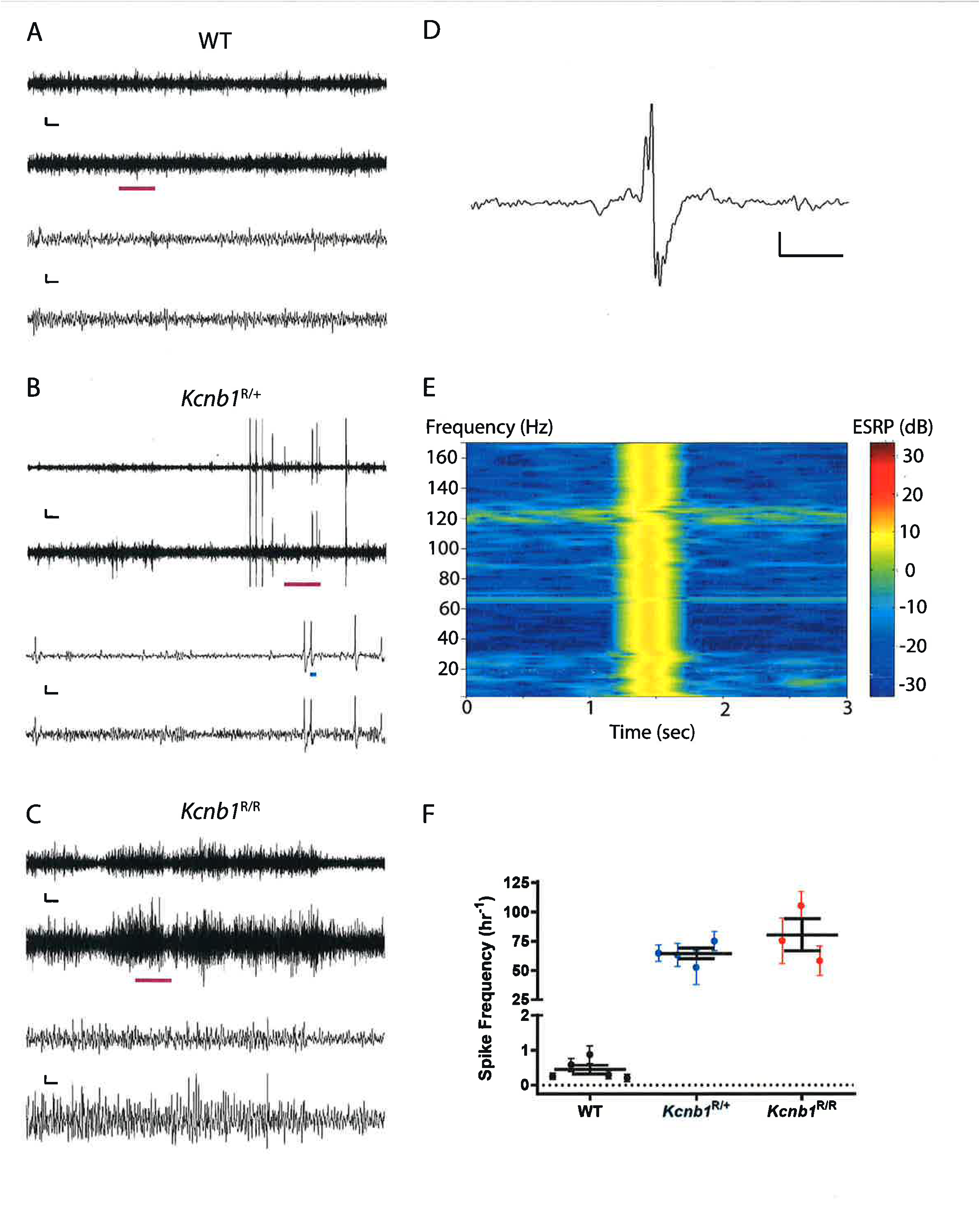
EEG abnormalities in *Kcnb1*^G379R^ mice. **A-C)** Representative EEG traces from WT, *Kcnb1*^R/+^, and *Kcnb1*^R/R^ mice, respectively. The top line corresponds to EEG1 and the second line corresponds to EEG2 in each set of traces. The top two lines display 5 minutes of EEG and the purple line denotes the 30 second region of expanded time base displayed below. The scale bar for the 5 minute time base are 75 µV and 10 sec and the scale bar for the 30 second segment are 75 µV and 1 sec, respectively. **D)** Example isolated spike and slow wave discharge corresponding to blue line in **B**. **E)** Power spectrum for spike and slow wave discharge in **D** showing elevated power in decibels across the 1-170 Hz frequency range at the time of the discharge. **F)** Quantification of isolated spike and slow wave discharges for each genotype over a 24 hour period and displayed as frequency per hour. The black lines and bars represent the mean ± SEM for all animals of that genotype and the individual symbols represent the mean for an individual animal ± SEM.

### Neurological abnormalities in *Kcnb1^G379R^* mice

A neurological exam was used to evaluate baseline neurological behavior in 3 week old WT, *Kcnb1*^R/+^ and *Kcnb1*^R/R^ mice, including analysis of muscle, spinocerebellar, sensory, neuropsychiatric and autonomic functions (Irwin, 1968). Neurological exam scores were affected by *Kcnb1* genotype, with higher scores indicating deficits. *Kcnb1*^R/+^ had an average overall exam score of 17.2 ± 0.4, that was elevated (p=0.0005) compared to the WT control score of 15.1 ± 0.4 (Figure 7A). *Kcnb1*^R/R^ mice were found to have the highest score, with an average of 19.8 ± 0.5 (Figure 7A) (p<0.0001 vs WT and *Kcnb1^R/+^*, respectively). Differences in the overall exam score were driven largely by differences in activity levels, escape behavior, and trunk curl (Supplemental Table S1). Furthermore, it was noted that upon transfer into the observation cage, 2 of 23 *Kcnb1*^R/R^ mice exhibited a tonic-clonic seizure characterized by forelimb clonus with rearing and falling.

**Figure 7.**
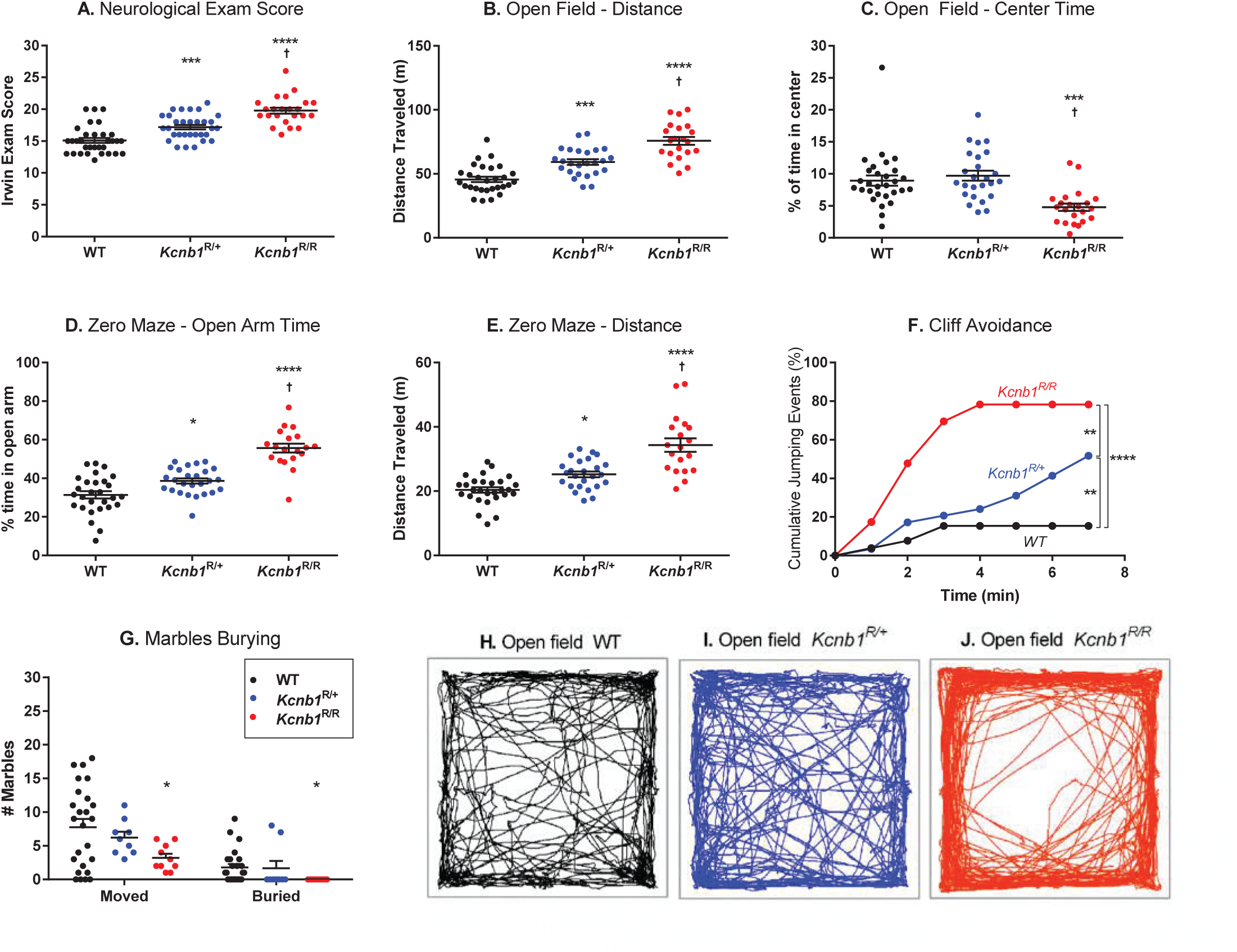
Neurological and neurobehavioral phenotypes in *Kcnb1^R/+^* and *Kcnb1^R/R^* mice. **A)** Modified Irwin neurological exam scores were affected by genotype (F(2,83) = 33.09, p<0.0001; one-way ANOVA). *Kcnb1^R/+^* mice (n=31) had an average score of 17.2 ± 0.4; higher compared to the WT (n=32) score of 15.1 ± 0.4 (***p = 0.0005, Tukey’s post-hoc comparisons). *Kcnb1^R/R^* mice (n=23) had the highest score, with an average of 19.8 ± 0.5, compared to WT and *Kcnb1^R/+^* mice (****,† p<0.0001, respectively; Tukey’s). **B & C)** Baseline activity was assessed using an open-field assay. **B)** There was a significant effect of genotype on total distance traveled (F(2,72) = 39.11, p<0.0001; one-way ANOVA). *Kcnb1^R/+^* mice (n=25) traveled an average distance of 59.3 ± 2.2 m, farther compared to WT (n=29) average distance of 45.6 ± 2.0 m (***p = 0.0002, Tukey’s). *Kcnb1^R/R^* mice (n=21) traveled an average distance of 75.8 ± 3.1 m, farther than both WT and *Kcnb1^R/+^* mice (****,† p<0.0001, respectively, Tukey’s). **C)** There was a significant effect of genotype on percent of time spent in the center of the open field (F(2,73) = 11.56, p<0.0001; one-way ANOVA). *Kcnb1^R/R^* mice spent 4.8 ± 0.8% of time in the center, less than both WT (***p = 0.0006, Tukey’s) and *Kcnb1^R/+^* mice (†p<0.0001, Tukey’s). **D & E)** Anxiety-related behavior was assessed using a zero-maze assay. **D)** Time spent in the open arms of the zero-maze was affected by genotype (F(2,70) = 42.07, p<0.0001; one-way ANOVA). *Kcnb1^R/+^* mice (n=26) spent an average of 38.6 ± 1.3% of the test time in open arms of the maze compared to WT (n=28), who averaged 31.4 ± 1.9% of time (*p = 0.0122, Tukey’s). *Kcnb1^R/R^* mice (n=19) spent 55.7 ± 2.3% of time in the open arms of the maze, more than both WT and *Kcnb1^R/+^* mice (****,†p< 0.0001, respectively, Tukey’s). **E)** There was a significant effect of genotype on total distance traveled in the zero maze (F(2,69) = 30.64, p<0.0001; one-way ANOVA). Total distance traveled was 34.3 ± 2.1 m for *Kcnb1^R/R^*, 25.2 ± 0.9 m for *Kcnb1^R/+^*, and 20.4 ± 0.8 m for WT (*p = 0.0125 and ****p<0.0001 vs WT, †p<0.0001 vs *Kcnb1*^R/+^, Tukey’s). **F)** Time to step or jump off the platform was evaluated in the cliff avoidance assay. *Kcnb1^R/+^* mice left the platform more frequently (≈52%, 15/29) compared to WT (≈15%, 4/26) (**p=0.0088, LogRank Mantel-Cox). *Kcnb1^R/R^* mice (≈78%, 18/23) left the platform more often than both WT and *Kcnb1^R/+^* mice (****p<0.0001, **p = 0.0052, respectively, LogRank Mantel-Cox). **G)** Marble burying was significantly affected by genotype (F(2,41) = 3.518, p = 0.0389; two-way repeated measures ANOVA). Relative to WT (n=25), *Kcnb1^R/+^* mice (n=9) rarely and *Kcnb1^R/R^* mice (n=10) never buried marbles (R/R vs WT, *p = 0.0336, Sidak’s). **H-J)** Representative examples of movement paths in the open-field assay for WT, *Kcnb1^R/+^* and *Kcnb1^R/R^* mice. Scatter plot data represents samples from individual mice, long horizontal lines represent the mean, and error bars represent SEM.

### *Kcnb1^G379R^* mice exhibit profound hyperactivity

*Kcnb1^G379R^* mice were easily distinguishable from WT littermates in home cages based on their elevated activity level. Baseline activity measured in an open field assay showed that *Kcnb1*^R/+^ mice traveled farther than WT controls, with average distances of 59.3 ± 2.2 m and 45.6 ± 2.0 m, respectively (p=0.0002) (Figure 7B). *Kcnb1*^R/R^ mice spent the majority of the session moving around the perimeter of the arena, traveling an average distance of 75.8 ± 3.1 m, more than both WT controls and *Kcnb1*^R/+^ mice (p<0.0001 and p<0.0001, respectively) (Figure 7B,C,J). *Kcnb1*^R/R^ mice spent <5% of their time in the center of the arena compared to WT and *Kcnb1*^R/+^ mice that spent ∼10% of the time in the center (Figure 7C, H, I).

### Impulsivity and diminished anxiety-related behaviors in *Kcnb1^G379R^* mice

Along with their noticeably elevated activity, *Kcnb1* mutants appeared to have altered impulsivity/anxiety-like behavior. To further assess these behavioral abnormalities, we used the zero-maze, cliff avoidance and marble burying assays. The zero-maze assay evaluated anxiety-related behavior of WT and *Kcnb1* mutants by comparing time spent in open versus closed arms. *Kcnb1*^R/+^ mice spent an average of 38.6 ± 1.3% of the test time in open arms compared to WT controls that averaged 31.4 ± 1.9% of time (p=0.0122) (Figure 7D). *Kcnb1*^R/R^ mice spent more than half their time (55.7 ± 2.3%) in the open arms of the maze, more than both WT and *Kcnb1*^R/+^ littermate mice (p< 0.0001, respectively) (Figure 7D). Distance traveled in the zero-maze differed between genotypes and mirrored the effects seen in open field distance traveled (Figure 7E).

The cliff avoidance assay takes advantage of the natural tendency of mice to avoid a potential fall from a height and is used to assess inattentive and impulsive behavior (Matsuoka et al., 2005; Yamashita et al., 2013). WT and *Kcnb1^G379R^* mice were individually placed on an elevated platform and monitored for jumping or stepping off the platform. WT mice rarely (4 of 26) left the platform (Figure 7F). *Kcnb1^R/+^* (≈52%; 15 of 29) and *Kcnb1^R/R^* (≈78%; 18 of 23) mice left the platform more frequently than WT controls (p=0.0088 and p<0.0001, respectively) (Figure 7F). In addition, there was an allele dosage effect with homozygous *Kcnb1*^R/R^ mice leaving the platform more often than heterozygous *Kcnb1*^R/+^ mice (p=0.0052).

The marble burying assay assessed attention, anxiety, and obsessive-compulsive related behaviors of WT and *Kcnb1* mutants. Relative to WT controls, *Kcnb1*^R/R^ mice showed little interaction with the marbles, moving few and failing to bury any (p=0.0336) (Figure 7G).

### *Kcnb1^G379R^* mice exhibit prolonged QT interval

To evaluate cardiac function in *Kcnb1^G379R^* mice, we performed electrocardiography (ECG) and echocardiography (ECHO) studies in *Kcnb1^G379R^* and WT mice (Figure 8). Heart rate-corrected QT interval (QT_C_) was prolonged in *Kcnb1^R/R^* relative to WT littermates, both at baseline (p=0.0292) (Figure 8B,C) and 6 minutes after isoproterenol administration (p=0.0002) (Figure 8B,D). Furthermore, *Kcnb1^R/R^* QT_C_ interval following isoproterenol injection was prolonged relative *Kcnb1^R/+^* (p=0.0202) (Figure 8B,D). There were no genotype-dependent differences detected in ejection fraction or fractional shortening determined by echocardiography (Supplemental Figure S1, Table S2), indicating the absence of contractile dysfunction at 10-14 weeks of age.

**Figure 8.**
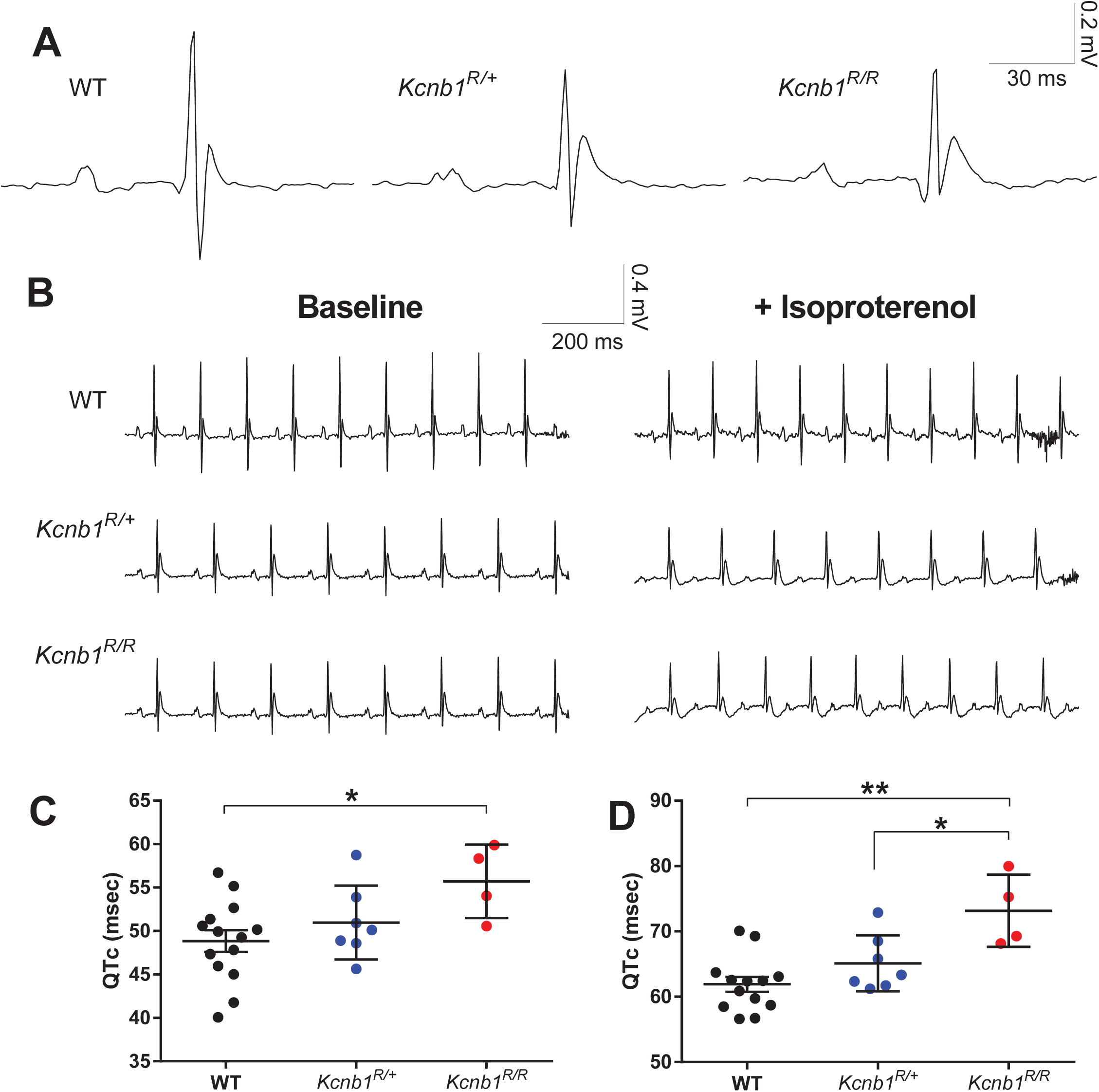
Cardiac QT interval is prolonged in *Kcnb1^R/R^* mice. Surface electrocardiograms (ECG) were recorded in 8-10 week old WT, *Kcnb1^R/+^* and *Kcnb1^R/R^* mice under anesthesia. Baseline recording was acquired for at least 4 minutes prior to adrenergic challenge with isoproterenol (1.5 mg/kg, IP) **A)** Representative recording of a single sinus beat at baseline in WT, *Kcnb1^R/+^* and *Kcnb1^R/R^* mice. **B)** Representative ECG traces from each genotype at baseline (left) and following isoproterenol administration (right). **C-D)** Both at baseline (C) and 6 minutes following isoproterenol administration (D), heart-rate corrected QT interval (QT_C_) was affected by genotype (F(2,21)=7.19; two-way repeated measures ANOVA). **C)** At baseline, *Kcnb1^R/R^* mice had a QT_C_ interval of 55.7 ± 2.1 msec, which was prolonged relative to the WT QT_C_ interval of 48.8 ± 1.2 msec (*p=0.0292, Sidak’s). **D)** Following isoproterenol (1.5 mg/kg, IP) administration, *Kcnb1^R/R^* mice had a QT_C_ interval of 73.2 ± 2.8 msec, which was prolonged relative to the WT and *Kcnb1^R/+^* QT_C_ intervals of 62.0 ± 1.2 and 65 ± 1.6 msec, respectively (**p=0.0002 and *p=0.0202, respectively; Sidak’s).

## Discussion

*KCNB1-*G379R was identified as a pathogenic *de novo* variant in a child with DEE. Electrophysiological studies of K_V_2.1 channels with the G379R variant in a heterologous expression system demonstrated lower potassium conductance relative to WT, loss of ion selectivity, and gain of a depolarizing inward cation conductance (Torkamani et al., 2014). In addition, mutant subunits exerted a dominant negative effect on potassium conductance when co-expressed with WT subunits (Torkamani et al., 2014). We extended *in vitro* characterization of the *KCNB1*-G379R variant by demonstrating failure of K_V_2.1-G379R to induce ER-PM junctions, as well as dominant negative effects on co-expressed WT K_V_2.1 subunits. This indicated that *KCNB1*-G379R results in deficits in both conducting and non-conducting K_V_2.1 functions. Introduction of this variant in *Kcnb1^G379R^* knock-in mice provides a new model of *KCNB1* associated developmental and epileptic encephalopathy. This model recapitulates several key features of the human disorder, including seizures, interictal epileptiform events on EEG, behavioral hyperactivity, impulsivity/inattention, and attenuated anxiety-related behavior.

The proband with the *KCNB1*-G379R *de novo* variant initially presented with infantile spasms at 8 months of age and later developed multiple seizure types including focal dyscognitive, atonic, and generalized tonic-clonic seizures (Torkamani et al., 2014). Seizures were difficult to treat and adequate control was not achieved with ACTH, topiramate, valproic acid, pyridoxine, or the ketogenic diet. EEG findings at different time points included hypsarrhythmia, diffuse polyspikes, diffuse polyspike and slow-waves, right temporal spike and waves, and left occipital spikes. Beyond seizures, there was also developmental delay and an autism spectrum disorder diagnosis consistent with atypical Rett syndrome, as well as borderline long QT syndrome (Srivastava et al., 2018; Torkamani et al., 2014). The *Kcnb1^G379R^* mouse model shares a number of these key phenotypes and will be a useful platform for evaluation of potential therapies for *KCNB1* associated DEE.

Electrographic and behavioral seizures in the mice shared features of seizures seen in the proband (Torkamani et al., 2014). The *Kcnb1^G379R^* mice exhibited multiple seizure types, including epileptiform events coinciding with behavioral arrest reminiscent of focal dyscognitive seizures, as well as generalized tonic-clonic seizures. Numerous EEG abnormalities were noted in *Kcnb1^G379R^* mice, including isolated spike and slow waves and recurrent runs of rhythmic slow spike and wave complexes. These events and features will be useful biomarkers to evaluate potential therapeutic strategies that can normalize the EEG.

Neurobehavioral abnormalities were prominent in the *Kcnb1^G379R^* mice and overlap with features reported in children with *KCNB1* encephalopathy (Bar et al., 2019; Calhoun et al., 2017; de Kovel et al., 2017; Marini et al., 2017; Srivastava et al., 2018; Thiffault et al., 2015; Torkamani et al., 2014). Attention-deficit/hyperactivity disorder or hyperactivity with inattention has been reported in numerous cases of *KCNB1* encephalopathy. *Kcnb1^G379R^* mice exhibit profound hyperactivity both in their home and novel environments, and inattention was suggested by failure to interact with marbles in the marble burying assay. In addition, lack of preference for the closed arms in the zero maze could be due to inattention to surroundings and/or reduced anxiety-like behavior. Failure of the cliff avoidance response in *Kcnb1^G379R^* mice may reflect inattention and/or elevated impulsivity, another behavioral problem reported for individuals with *KCNB1* encephalopathy. Although visual deficits could explain the observed inattention, there was no deficit in visual placing in *Kcnb1^G379R^* mice and prior characterization of *Kcnb1^-/-^* mice included assessment of vision, which detected no impairment (Speca et al., 2014). Autism spectrum disorder has been reported in more than half of *KCNB1* encephalopathy cases, including *KCNB1*-G379R. Repetitive jumping was another prominent behavior observed in the *Kcnb1^G379R^* mice in their home cages and upon transfer to a novel environment. This perseverative behavior was also previously reported in the *Kcnb1^-/-^* mice and may reflect repetitive movements seen in *KCNB1* DEE patients with autism spectrum disorder (Speca et al., 2014).

Previous work demonstrated that global deletion of *Kcnb1* in mice resulted in pronounced hyperactivity, reduced anxiety-like behavior, and increased susceptibility to flurothyl and pilocarpine-induced seizures (Speca et al., 2014). The neurobehavioral phenotypes of *Kcnb1^R/R^* mice and *Kcnb1^-/-^* mice were largely similar on the same C57BL/6J background strain. However, the more prominent seizure phenotypes in *Kcnb1^G379R^* mice suggest the mutation is not simply a loss-of-function allele. Spontaneous seizure activity was reported in ≈10% of homozygous *Kcnb1*^-/-^ mice during the course of routine handling (Speca et al., 2014). This is similar to the frequency of spontaneous seizure activity observed in heterozygous *Kcnb1^R/+^* mice, while seizures were observed in almost half of homozygous *Kcnb1^R/R^* mice between 6 and 12 weeks of age. *In vitro* functional studies of *KCNB1*-G379R showed change of function effects including loss of voltage-sensitivity and altered ion-selectivity, as well as dominant negative effects when co-expressed with WT subunits (Torkamani et al., 2014). In addition to alterations in the biophysical properties of K_V_2.1 channels, the G379R mutation also appeared to exert a dominant negative influence on the expression and subcellular localization of WT K_V_2.1, K_V_2.2 and AMIGO-1 subunits. Immunohistochemistry and immunoblotting experiments revealed that both K_V_2.2 and AMIGO-1 immunoreactivity was substantially lower in sections and whole brain membrane preparations from *Kcnb1^R/R^* mice. K_V_2.1 and K_V_2.2 form heteromeric channels, with most complexes containing AMIGO-1 auxiliary subunits (Bishop et al., 2018; Kihira et al., 2010; Peltola et al., 2011a). We hypothesize that the lower K_V_2.2 and AMIGO-1 immunoreactivity is due to their altered subcellular localization, as both proteins were diffusely distributed throughout the ER rather than clustered at the PM (for a full discussion see (Bishop et al., 2018)). The expression and localization alterations resulting from the *Kcnb1^G379R^* mutant may contribute to the more severe seizure phenotypes observed in *Kcnb1^G379R^* mice as compared to *Kcnb1^-/-^* null mice. Future studies will investigate the neurophysiological basis of seizure phenotypes in *Kcnb1^G379R^* mice.

Beyond neurological phenotypes, there have been anecdotal reports of borderline long QT syndrome in patients with *KCNB1* associated encephalopathy (kcnb1.org). Additionally, *KCNB1* is highly expressed in mouse cardiac tissue and synonymous SNPs in *KCNB1* have been associated with long QT syndrome in humans (Iwasa et al., 2001). Therefore, we performed preliminary evaluation of cardiac phenotypes using ECG and ECHO. Homozygous *Kcnb1^R/R^* mice had prolonged QT_C_ interval relative to WT at baseline and following an isoproterenol challenge. K_V_2.1 has been shown to contribute to the I_Kslow_ component of the mouse cardiac action potential, and a transgenic K_V_2.1 dominant-negative mutation (N216) was shown to attenuate I_Kslow_ resulting in action potential and QT prolongation (Xu et al., 1999). There was no evidence of structural abnormalities or contractile dysfunction by ECHO (Supplemental Table S2). Our results suggest that *KCNB1* variants could conceivably contribute to spontaneous or induced cardiac arrhythmia (e.g., exposure to proarrythmic drugs), although future studies will be required to further characterize the arrhythmogenic potential of *Kcnb1^G379R^*. Interestingly, mice with homozygous deletion of AMIGO-1, an auxiliary subunit of the K_V_2.1 channel complex (Peltola et al., 2011b), also have a prolonged QT_C_ interval (Dickinson et al., 2016). This provides additional evidence associating the K_V_2.1 channel complex with QT_C_ prolongation.

In summary, we developed a novel knock-in mouse model of human DEE caused by a missense mutation in *Kcnb1* that disrupts both conducting and non-conducting functions of K_V_2.1 channels. The *Kcnb1^G379R^* mouse model will be valuable for defining the molecular and neurophysiological consequences of *Kcnb1* mutation, understanding disease pathophysiology, and evaluating response to therapeutic interventions.

## Acknowledgements

We thank Alexandra Huffman and Nicole J. Zachwieja for technical assistance. The genetically engineered mice were generated with the assistance of Lynn Doglio in Northwestern University Transgenic and Targeted Mutagenesis Laboratory. Echocardiography was performed by Veronica Ramirez in the Cardiovascular Phenotyping Core of the Feinberg Cardiovascular and Renal Research Institute. Sunita Misra was supported by a Lurie Children’s Hospital Pediatric Physician Scientist Research Award and Lisa Wren was supported by an American Heart Association Predoctoral Fellowship. This work was supported by the National Institutes of Health grants R01 NS053792 (JAK) and U54 NS108874.

## Declaration of interest

None.

## Notes

#### Summary of Updates

Figure 2 revised to correct miscoloring of epitope label text on panels A-D. Figure 3 revised to add lane labels to panels D and F.

