## Supplement for "Epilepsy and neurobehavioral abnormalities in mice with a *KCNB1* pathogenic variant that alters conducting and non-conducting functions of K_V_2.1"

for

**List of Supplemental Materials**

1. **Supplemental Materials and Methods.**
2. **Supplemental Table S1.** Neurological exam scores from wild-type (WT), *Kcnb1*$^{R/+}$ and *Kcnb1*$^{R/R}$ mice.
3. **Supplemental Table S2.** Echocardiogram data from wild-type (WT), *Kcnb1*$^{R/+}$ and *Kcnb1*$^{R/R}$ mice.
4. **Supplemental Figure S1.** *Kcnb1*$^{R/+}$ and *Kcnb1*$^{R/R}$ mice do not differ from wild-type (WT) by echocardiography.
5. **Supplemental Video 1.** Handling-induced behavioral seizure in a *Kcnb1*$^{R/R}$ mouse at 2-3 months of age. <Suppl Video S1 Kcnb1-RR Handling.mp4>
6. **Supplemental Video 2.** Paroxysmal event on EEG associated with immobility in a *Kcnb1*$^{R/R}$ mouse at 2-3 months of age. <Suppl Video S2 Kcnb1-RR Event.mp4>

**Supplemental Materials and Methods**

***HEK293T cell culture, immunocytochemistry, and epifluorescence and TIRF imaging***.

HEK293T cells were maintained in Dulbecco's modified Eagle's medium supplemented with 10% Fetal Clone III (HyClone Cat# SH30109.03), 1% penicillin/streptomycin, and 1x GlutaMAX (ThermoFisher Cat# 35050061) in a humidified incubator at 37 °C and 5% CO2. Plasmid constructs used include wild-type rat Kv2.1 (rKv2.1) in pRBG4 (Shi et al., 1994), rat Kv2.1 S586A (rKv2.1 S586A) in pCGN (Lim et al., 2000), HA-tagged wild-type human Kv2.1 (hKv2.1) (Kang et al, 2019; modified from Addgene #131707), and HA-tagged human Kv2.1 with the G379R mutation (hKv2.1 G379R)(Kang et al, 2019; modified from Addgene #131707). For transfections, cells were split to 15% confluence on number 1.5 glass coverslips coated with poly-L lysine, then transiently transfected using Lipofectamine 2000 following the manufacturer's protocol within 18 hours of plating. Cells were transiently transfected in DMEM without supplements, then returned to regular growth media 4 hours after transfection. Cells were utilized 40-48 hours post-transfection. For experiments involving immunocytochemistry, fixation was performed as previously described (Dickson et al., 2016; Kirmiz et al., 2018). Briefly, HEK293T cells were fixed in 3.2% formaldehyde (freshly prepared from paraformaldehyde, Sigma Cat# 158127) and 0.1% glutaraldehyde (Ted Pella, Inc., Cat# 18426) for 30 minutes and room temperature, washed 3, 5 minutes in PBS and quenched with 1% sodium borohydride in PBS for 15 minutes at room temperature. Cells were blocked and permeabilized in 4% non-fat milk powder in TBS containing 0.1% Triton-X 100. Primary antibody incubation was performed in blocking solution for 1 hour at room temperature. Primary antibodies used (see Table 2 for details) were mouse monoclonal anti-HA epitope tag antibody 12CA5 (RRID:AB_ 2532070, pure, 5 μg/mL) to detect exogenous hKv2.1, rKv2.1 and hKv2.1 G379R, and mouse monoclonal VAPA antibody N479/24 (RRID:AB_ 2722709, tissue culture supernatant, 1:5) to detect endogenous VAPA (Kirmiz et al., 2018). Following primary antibody incubation and 3, 5 minute washes in blocking solution at room temperature, coverslips were immunolabeled in blocking solution with mouse IgG subclass-specific Alexa Fluor-conjugated goat anti-mouse IgG subclass-specific (Manning et al., 2012) secondary antibodies (all secondary antibodies from ThermoFisher) at a 1:1500 dilution, and Hoechst 33258 (ThermoFisher Cat# H1399) at 200 ng/mL for one hour, washed 3, 5 min in PBS, and mounted in phosphate buffered saline onto glass depression slides.

Epifluorescence and TIRF imaging of fixed cells and image analysis was performed essentially as described (Bishop et al., 2018; Kirmiz et al., 2018). Images were obtained with an Andor iXon EMCCD camera installed on a TIRF/widefield equipped Nikon Eclipse Ti microscope using a Nikon LUA4 laser launch with 405, 488, 561, and 647 nm lasers and a 100x PlanApo TIRF/1.49 NA objective run with NIS Elements software (Nikon). Images were collected within NIS Elements as ND2 images.

Colocalization analyses were performed within Nikon NIS Elements using ND2 files. An ROI was drawn within a cell and Pearson's correlation coefficient (PCC) was collected, as previously described (Kirmiz et al., 2018). Group PCC values were compared using one-way ANOVA with Tukey's post-hoc comparisons (GraphPad Prism). Measurements of structure sizes were quantified automatically within Fiji previously described (Dickson et al., 2016). ND2 files collected in TIRF were imported to Fiji, background subtracted, converted into an 8-bit image, and automatically converted into a binary mask using auto local thresholding. An ROI with identical dimensions was drawn within each cell analyzed. The number of individual ER-PM junctions, average ER-PM junction size, and percentage PM occupancy were quantified automatically using the "analyze particles" function in Fiji, as previously described (Kirmiz et al., 2018). Signals smaller than 0.04 $\mu m^2$ were excluded from this analysis. Puncta size values were compared using one-way ANOVA with Tukey's post-hoc comparisons (GraphPad Prism).

***Immunohistochemistry.*** Animals were deeply anesthetized with pentobarbital and transcardially perfused with 4% formaldehyde prepared from paraformaldehyde, in 0.1 M sodium phosphate buffer pH 7.4 (0.1 M PB). Sagittal brain sections (30 µm thick) were prepared and immunolabeled using free-floating methods as previously described (Palacio et al., 2017; Rhodes et al., 2004; Speca et al., 2014). Free floating sections were blocked and permeabilized with 10% goat serum, 0.3% Triton X-100 in 0.1 M PB for 1 hour at room temperature (RT) and incubated overnight at 4°C in primary antibodies (Table 2). After 4, 5 min washes, sections were exposed to species- and/or mouse IgG subclass-specific Alexa-conjugated fluorescent secondary antibodies (Invitrogen) and Hoechst 33258 DNA stain diluted in vehicle for 1 hour at RT. After 2, 5 min washes in 0.1M PB and a 5 min wash in 0.05M PB, sections were mounted and air dried onto gelatin-coated microscope slides, treated with 0.05% Sudan Black (EM Sciences) in 70% ethanol for 2 min (Schnell et al., 1999), extensively washed in water, and mounted with Prolong Gold (ThermoFisher Cat # P36930). Images were taken using the same exposure time to compare the signal intensity directly, using an AxioCam HRm high-resolution CCD camera installed on an Axioskop M2 or AxioObserver Z1 microscope with 63x, 1.3 numerical aperture (NA) lens or 20x, 0.8 NA lens, and an ApoTome coupled to Axiovision software, version 4.8.2.0 (Zeiss, Oberkochen, Germany). Images were identically processed in Photoshop to maintain consistency between samples. Labeling intensity within *stratum pyramidale* of hippocampal CA1 was measured using a rectangular region of interest (ROI) of 39 µm x 164 µm. To maintain consistency between samples, ROIs were obtained from a region within CA1 near the border of CA1 and CA2. Data points reflect the mean pixel intensity values of this ~6400 $\mu m^2$ ROI. Values from multiple immunolabels and of Hoechst dye were simultaneously measured from the same ROI. Background levels for individual labels were measured from no primary controls and mathematically subtracted from ROI values. Data is represented as mean ± SEM. Group values were compared using one-way ANOVA with Tukey's post-hoc comparisons (GraphPad Prism).

***Echocardiography.*** Male and female WT, *Kcnb1*$^{R/+}$ and *Kcnb1*$^{R/R}$ mice were imaged between P74 and P98. Mice were anesthetized with 2-4% isoflurane while on a warming platform with a rectal probe to ensure body temperature was maintained. Imaging duration was 30-40 minutes per mouse. Echocardiography was performed using Visualsonics Vevo770 (FujiFilm, Toronto, Canada). The following parameters were measured: left ventricular end-diastolic dimension (LVEDD), left ventricular end-systolic dimension (LVESD), percent fractional shortening (%FS), calculated as (LVEDD-LVESD)/LVEDD×100, septal thickness (SEPth), posterior wall thickness(PWth), heart rate corrected mean velocity of circumferential shortening (mVcfc), calculated as %FS / ejection time × the square root of the R-R interval. Estimated echocardiographic LV mass (in mg) was calculated as ((LVEDD + SEPth + PWth)3 - LVEDD3) × 1.055, where 1.055 (mg/mm3) is the density of myocardium. Statistical comparison between groups was made using ANOVA with Tukey's post-hoc tests.

**Supplemental Table S1**. Neurological exam scores from wild-type (WT), *Kcnb1*$^{R/+}$ and *Kcnb1*$^{R/R}$ mice. Individual parameters were compared to WT using multiplicity-adjusted T-tests with a false-discovery rate of 1%. Discoveries are indicated in bold.

| | WT (n=32) | | *Kcnb1*$^{R/+}$ (n=31) | | | *Kcnb1*$^{R/R}$ (n=23) | | |
|---|---|---|---|---|---|---|---|---|
| Parameter | Mean | SD | Mean | SD | P-value* vs WT | Mean | SD | P-value* vs WT |
| Transfer Behavior | 3.8 | 0.5 | 4 | 0.3 | 0.0488 | 4.3 | 0.5 | **0.0002** |
| Spontaneous Activity | 2.4 | 0.6 | 2.9 | 0.6 | **<0.0001** | 3.8 | 0.5 | **<0.0001** |
| Tremor | 0.1 | 0.4 | 0.4 | 0.6 | 0.0188 | 0.3 | 0.5 | 0.0665 |
| Gait | 0.0 | 0.0 | 0 | 0.2 | 0.7722 | 0.4 | 0.5 | **0.0004** |
| Pelvic Elevation | 2.0 | 0.0 | 2 | 0 | 1.0000 | 2 | 0 | 1.0000 |
| Tail Elevation | 1.0 | 0.0 | 1 | 0 | 1.0000 | 1 | 0 | 1.0000 |
| Palpebral Closure | 0.0 | 0.0 | 0 | 0 | 1.0000 | 0 | 0.2 | 0.7200 |
| Piloerection | 0.0 | 0.0 | 0 | 0 | 1.0000 | 0.1 | 0.3 | 0.4735 |
| Escape Behavior | 0.7 | 0.7 | 1.3 | 0.6 | **<0.0001** | 1.8 | 0.6 | **<0.0001** |
| Jumping | 0.3 | 0.5 | 0.6 | 0.5 | 0.0071 | 0.5 | 0.5 | 0.1720 |
| Air Puff | 2.4 | 0.9 | 2.7 | 0.8 | 0.0068 | 3.5 | 0.6 | **<0.0001** |
| Trunk Curl | 0.8 | 0.4 | 0.4 | 0.5 | **0.0012** | 0 | 0 | **<0.0001** |
| Limb Grasping | 0.3 | 0.5 | 0.2 | 0.4 | 0.2819 | 0.1 | 0.3 | 0.1095 |
| Preyer Reflex | 0.7 | 0.5 | 0.9 | 0.7 | 0.0981 | 1.3 | 0.8 | **<0.0001** |
| Salivation | 0.0 | 0.0 | 0 | 0 | 1.0000 | 0 | 0 | 1.0000 |
| Provoked Biting | 0.6 | 0.5 | 0.8 | 0.4 | 0.0577 | 0.7 | 0.5 | 0.4598 |

*False-discovery rate adjusted p-values (q=0.01).

**Supplemental Table S2.** Echocardiogram data from wild-type (WT), *Kcnb1*$^{R/+}$ *and Kcnb1*$^{R/R}$ *mice.*

### Ejection Fraction

| Genotype | Sex | HR | Temp | *Diastolic* | | *Systolic* | | EF % |
|---|---|---|---|---|---|---|---|---|
| | | | | Area $mm^2$ | Volume ul | Area $mm^2$ | Volume ul | |
| WT | F | 521 | 39.1 | 21.05 | 57.22 | 10.8 | 21.16 | 63.02 |
| WT | F | 536 | 37.7 | 17.01 | 41.05 | 8.83 | 12.55 | 69.43 |
| WT | F | 521 | 39.2 | 19.51 | 50.17 | 10.75 | 17.58 | 64.96 |
| WT | M | 523 | 38.3 | 21.16 | 61.99 | 11.91 | 22.44 | 63.80 |
| WT | M | 537 | 36.7 | 14.74 | 34.49 | 7.65 | 11.45 | 66.80 |
| *Kcnb1*$^{R/+}$ | F | 483 | 38.3 | 18.22 | 45.64 | 10.06 | 16.96 | 62.84 |
| *Kcnb1*$^{R/+}$ | F | 531 | 37.9 | 16.09 | 40.45 | 8.14 | 12.03 | 70.26 |
| *Kcnb1*$^{R/+}$ | M | 527 | 38.4 | 17.2 | 40.11 | 10.04 | 16.69 | 58.39 |
| *Kcnb1*$^{R/+}$ | M | 518 | 38 | 15.61 | 40.72 | 7.24 | 11.89 | 70.80 |
| *Kcnb1*$^{R/+}$ | M | 541 | 38.1 | 15.36 | 31.66 | 8.33 | 11.98 | 62.16 |
| *Kcnb1*$^{R/R}$ | F | 465 | 37.7 | 13.91 | 31.62 | 6.2 | 9.76 | 69.13 |
| *Kcnb1*$^{R/R}$ | M | 512 | 38 | 20.8 | 56 | 9.61 | 17.3 | 69.11 |
| *Kcnb1*$^{R/R}$ | M | 517 | 37.5 | 21.22 | 57.91 | 9.74 | 16.43 | 71.63 |
| *Kcnb1*$^{R/R}$ | M | 553 | 38.5 | 20.89 | 56.78 | 10.52 | 18.35 | 67.68 |
| *Kcnb1*$^{R/R}$ | M | 552 | 38.3 | 16.02 | 38.61 | 9.24 | 16.39 | 57.55 |

### Fractional Shortening

| Genotype | Sex | HR | Temp | *Diastolic* | *Systolic* | FS % |
|---|---|---|---|---|---|---|
| | | | | DD | SD | |
| WT | F | 516 | 38.8 | 3.79 | 2.35 | 37.99 |
| WT | F | 532 | 37.1 | 3.27 | 1.8 | 44.95 |
| WT | F | 521 | 38 | 3.41 | 1.88 | 44.87 |
| WT | M | 523 | 38.3 | 4.08 | 2.57 | 37.01 |
| WT | M | 542 | 37.6 | 3.56 | 2.01 | 43.54 |
| *Kcnb1*$^{R/+}$ | F | n/a | n/a | 3.57 | 2.31 | 35.29 |
| *Kcnb1*$^{R/+}$ | F | 521 | 37.3 | 3.52 | 1.95 | 44.60 |
| *Kcnb1*$^{R/+}$ | M | 532 | 38.1 | 3.65 | 2.51 | 31.23 |
| *Kcnb1*$^{R/+}$ | M | 521 | 38 | 3.49 | 1.92 | 44.99 |
| *Kcnb1*$^{R/+}$ | M | 538 | 38.1 | 3.35 | 2.3 | 31.34 |
| *Kcnb1*$^{R/R}$ | F | 497 | 37 | 2.92 | 1.82 | 37.67 |
| *Kcnb1*$^{R/R}$ | M | 516 | 38.2 | 3.85 | 2.61 | 32.21 |
| *Kcnb1*$^{R/R}$ | M | 516 | 37.5 | 3.87 | 2.48 | 35.92 |
| *Kcnb1*$^{R/R}$ | M | 554 | 38.5 | 3.64 | 2.42 | 33.52 |
| *Kcnb1*$^{R/R}$ | M | 545 | 38.5 | 3.56 | 1.89 | 46.91 |

**Supplemental Figure S1.**

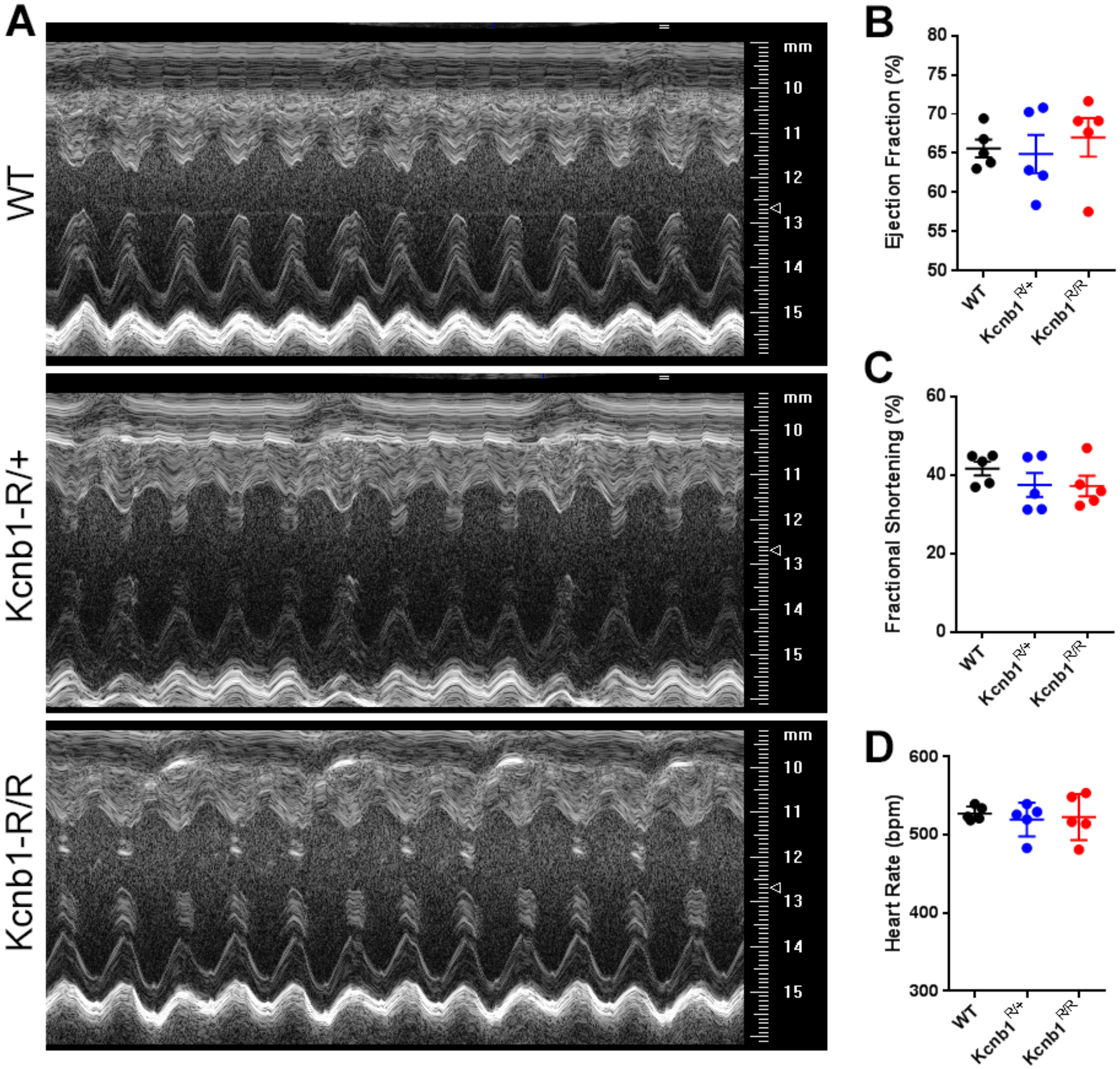


**Supplemental Figure S1**. *Kcnb1*$^{R/+}$ and *Kcnb1*$^{R/R}$ mice do not differ from wild-type (WT) by echocardiography. **A)**. Representative short-axis M-mode echocardiography images. B) Ejection fraction was not different between groups. C) Fractional shortening was not different between genotypes. D) Average heart rate did not differ between genotypes. Average ± SEM values are shown. Statistical comparison between groups was made using ANOVA with Tukey's post-hoc comparisons. Detailed underlying data for B-D are shown in Table S2.

**Supplemental Video S1**. Handling-induced generalized tonic-clonic seizure in a *Kcnb1*$^{R/R}$ mouse.

**Supplemental Video S2**. Slow spike and wave event in a *Kcnb1*$^{R/R}$ mouse coinciding with behavioral arrest.
